# Targeting BET Proteins downregulates miR-33a to promote synergy with PIM inhibitors in CMML

**DOI:** 10.1101/2022.11.01.514753

**Authors:** Christopher T. Letson, Maria E. Balasis, Hannah Newman, Moritz Binder, Alexis Vedder, Fumi Kinose, Markus Ball, Traci Kruer, Ariel Quintana-Gonzalez, Terra L. Lasho, Christy M. Finke, Luciana L. Almada, Jennifer M. Grants, Guolin Zhang, Martin E. Fernandez-Zapico, Alexandre Gaspar-Maia, Jeffrey Lancet, Rami Komrokji, Eric Haura, Gary W. Reuther, Aly Karsan, Uwe Rix, Mrinal M. Patnaik, Eric Padron

## Abstract

Preclinical studies in myeloid neoplasms have demonstrated efficacy of Bromodomain and Extra-Terminal protein inhibitors (BETi). However, BETi demonstrate poor single agent activity in clinical trials. Several studies suggest that combination with other anti-cancer inhibitors may enhance the efficacy of BETi. To nominate BETi combination therapies for myeloid neoplasms, we used a chemical screen with therapies currently in clinical cancer development. We identified PIM inhibitors (PIMi) as therapeutically synergistic with BETi in myeloid leukemia models. Mechanistically, we show that PIM kinase is increased after BETi treatment, and that PIM kinase upregulation is sufficient to induce resistance to BETi and sensitize cells to PIMi. Further, we demonstrate that miR-33a downregulation is the underlying mechanism driving PIM1 upregulation. We also show that GM-CSF hypersensitivity, a hallmark of chronic myelomonocytic leukemia (CMML), represents a molecular signature for sensitivity to combination therapy and credential this using patient-derived xenografts supporting the clinical investigation of this combination.

## Introduction

Mutations in genes governing epigenetic regulation are the most common alterations in myeloid malignancies. While clinically heterogenous, mutations in epigenetic regulators such as *TET2* and *DNMT3a* are predominantly early molecular events associated with disease initiation^1,2^. In a large subset of cases, secondary mutations in genes encoding signal transduction proteins such as *N/K-RAS* or *FLT3* result in leukemic transformation of clones harboring pre-existing epigenetic pathway mutations in both preclinical and clinical specimens^3,4^. The frequency and clonal composition of mutations in genes that alter the epigenome have suggested that therapies aimed at targeting this pathway may be therapeutically attractive. However, while several therapies have been tested across a variety of myeloid neoplasms, only 5-azacitidine has been clinically demonstrated to alter the natural history of myelodysplastic syndromes. Therefore, there exists a critical need to identify therapies that capitalize on the epigenetic dysregulation that molecularly hallmarks these cancers.

The development of potent inhibitors that bind to the BET family of proteins by mimicking acetylated lysine and occupying tandem bromodomains conserved among BET proteins represents a powerful epigenetic therapeutic approach that has been clinically tested in myeloid malignancies^5^. Occupation of tandem bromodomains prevents BET proteins from binding acetylated lysine resulting in widespread downregulation of gene expression, particularly those gene expression programs governed by super enhancers, such as NFkB signaling in myeloproliferative diseases^6,7^. Preclinical models of myeloid malignancies have suggested that BETi may be highly effective against a variety of hematological malignancies^8-10^. However, early phase BET inhibitor clinical trials have demonstrated minimal clinical efficacy with the potential for significant side effects associated with long term treatment^11^. Additionally, several resistance mechanisms to BET inhibitors have been identified such as Wnt pathway upregulation^12-14^, BRD4 hyperphosphorylation^15^, or transcriptional reprograming leading to alternate kinome dependencies^16^. Given the alterations in both epigenetic and signaling mutations seen in myeloid malignancies and the recent preclinical and clinical efficacy observed with combining BETi and JAKi in myeloproliferative neoplasms, we hypothesized that novel combinations of BETi and kinase inhibitors may represent an effective therapeutic strategy for myeloid malignancies^17^.

To test this hypothesis, we performed a broad in-house compound screen and identified PIMi as a potential synergistic combination with BETi. We validated this *in vitro* and *in vivo* utilizing human leukemia cells, primary patient material, and isogenic BETi persistent cells. Mechanistically, BETi dependent PIM1 overexpression was observed in cell lines vulnerable to the combination therapy and PIM1 overexpression alone was sufficient to both induce resistance to BETi and increase sensitivity to PIMi. We additionally demonstrate that BETi dependent PIM upregulation was secondary to downregulation of miR-33a, known post-transcriptional regulators of PIM1 via global repression of miRNA biogenesis. Last, we find that sensitivity to this combination is associated with cytokine dependent transcriptional priming of PIM1 and validate this in human specimens of chronic myelomonocytic leukemia (CMML) patients which are transcriptionally primed at the PIM1 loci and vulnerable to this therapeutic approach.

## Results

### Clinically relevant kinase inhibitor screen identifies BET and PIM inhibitors as synergistic in models of myeloid malignancies

In order to test potential synergies between BET inhibitors and inhibitors in clinical development, we utilized an in-house targeted chemical screen of 300 compounds which are FDA approved or in clinical cancer development (Supplemental table 1)^18^. U937 and SKM1 cells were incubated with the IC_20_ (U937:155nM, SKM1: 30nM) of the BET inhibitor INCB054329 and two doses (0.5μM and 2.5μM) of each library compound. Cell viability was evaluated 72hrs post-treatment using CellTitre-Glo. Combinations with a drug - base/drug + base ratio greater than 2 were chosen for further consideration as previously described^18^. As expected, known synergies with JAK, HDAC, CDK, MEK, and PI3K inhibitors were found supporting the validity of our chemical screen to identify clinically relevant BETi combinations^19-28^. After previously published interactions were filtered out (Supplemental Table 2), the only combination with a drug - base/drug + base ratio greater than 2 was with SGI-1776, a pan-PIM inhibitor (Fig. 1A). To validate therapeutic synergy between these BET and PIM inhibitors, we repeated the experiment in three human myeloid cell lines (U937, TF1 and SKM1) with 7 doses of INCB054329 and either pan-PIM inhibitors SGI-1776 or INCB053914. In all lines, and in both BET/PIM inhibitor combinations, *in vitro* synergy was observed consistent with our initial compound screen (Fig. 1B). Importantly, synergy was only evident in the low dose PIMi chemical screen and enhanced in most models when testing low doses of both BETi and PIMi (Fig 1B, blue circle).

**Figure 1.**
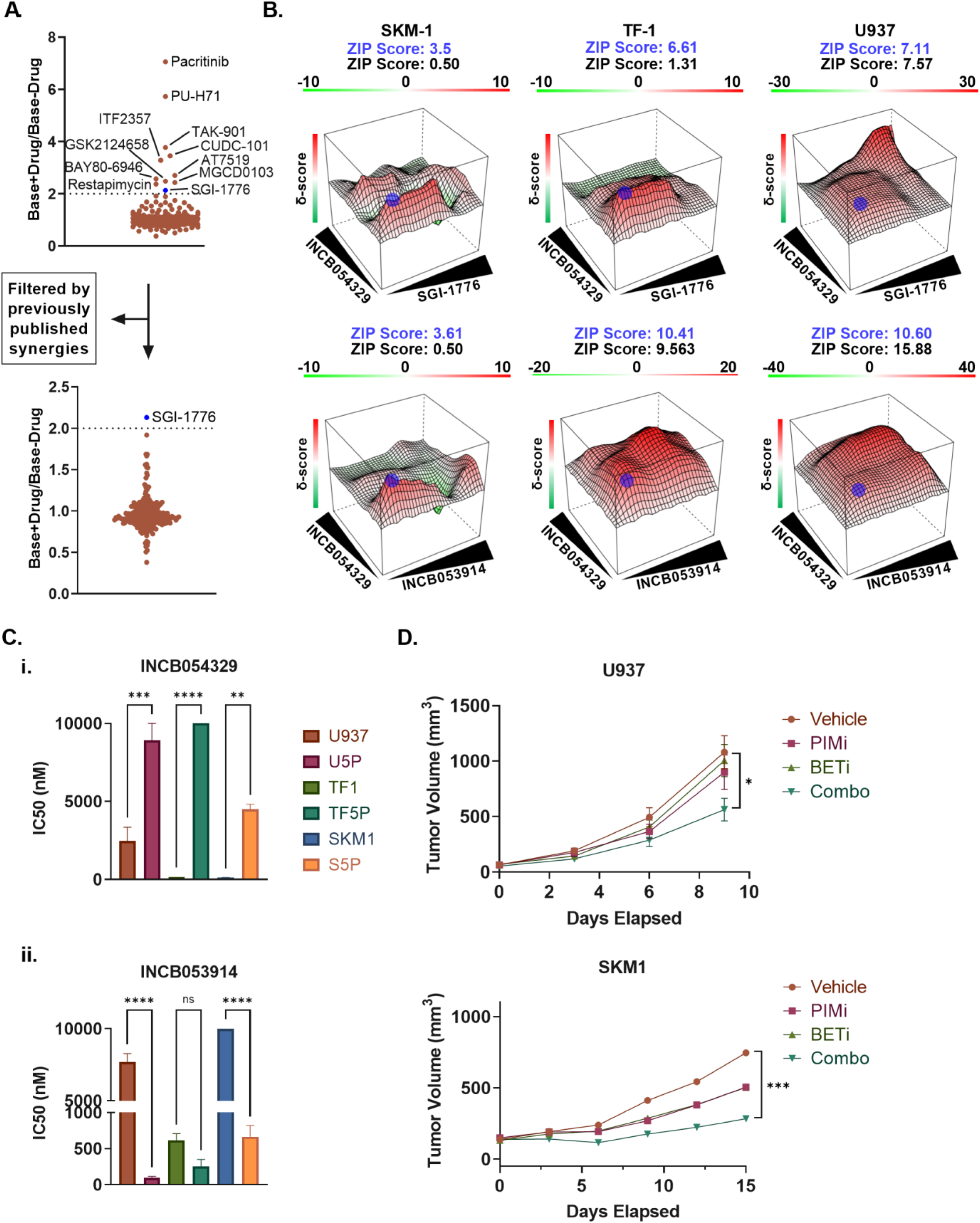
BETi and PIMi are synergistic in cell line models of CMML: (A)Results of kinase screen performed in U937 and SKM1 cells. Top: Ratio of base drug +/- experimental compounds for all targets. Bottom: Targets filtered by previously published research. (B) ZIP synergy plots generated by SynergyFinder of U937, SKM1 and TF1 cell lines; red indicates synergy, green indicates antagonism. Cell lines were treated with 7 increasing doses of both BETi and PIMi for 72hrs. (C) IC_50_ of parental cell lines and their persistent counterparts treated with either BETi(i) or PIMi(ii) for 72hrs. (D)Tumor size calculations of mice subcutaneously injected with either U937 or SKM1 cells and treated with BETi(INCB057643), PIMi(INCB053914) or combo (U937: Vehicle and BET n=10, PIMi n=8 Combo n=9; SKM1: Vehicle, BETi, PIMi n=9, Combo n=8). Mice were treated for 2 weeks unless tumors showed signs of ulceration. Significance was determined by comparing the Area Under the Curve (AUC). *=p<.05, ***=p<.0005.

Rapid persistence to BETi has been shown to occur through various mechanisms in both leukemia and solid tumors, likely responsible for its limited clinically efficacy as a single agent^12-16^. To determine whether PIMi could overcome persistence to BETi, we generated 3 BET persistent human leukemia cell lines. We then compared the IC_50_ of PIMi to that of the parental cell lines tested. Persistence was achieved by daily treatment of cell lines with 500nM INCB054329 or 300nM INCB054329 for SKM1 cells (Fig. 1Ci). At 60 days, all three cell lines demonstrated an increase in PIMi sensitivity compared to their parental counterparts, particularly in the human monocytic leukemia cell line U937 (Fig. 1Cii). To determine whether the observed *in vitro* synergy was present *in vivo*, heterotopic tumors were established in NSG-S mice^29^ with either U937 (n=10/group) or SKM1 cells (n=10/group). After tumors reached between 100 and 150mm^3^, drug treatment was started with 10 mpk INCB057643 and 30 mpk INCB053914 via oral gavage either as single agent or in combination and continued for 2 weeks, with tumor measurements occurring twice per week and at endpoint. These experiments identified a statistically significant decrease in tumor volume utilizing both cell line models with combination treatment suggesting that this combination strategy may be synergistic *in vivo (*U937: Vehicle AUC = 3778, Combo AUC = 2148, p<.05. SKM1: Vehicle AUC = 5483, Combo AUC = 2722, p<.0005. Figure 1D).

### PIM kinases are upregulated in response to BETi in a subset of leukemia cell lines and correlate to PIMi sensitivity

PIM proteins are serine/threonine kinases with a short half-life that do not require post-translational modifications for their activation and therefore, their activity is primarily transcriptionally and post-translationally mediated^30,31^. Given the profound effects that BET inhibitors exert on the transcriptome, we first sought to examine the effect of BET inhibition on RNA and protein expression of PIM kinases. Interestingly, PIM kinase protein and RNA expression of cells treated with BETi after 24 hrs revealed a significant increase in expression of PIM kinases. This increase was highest in BETi persistent cells where significant increases in PIM1 and PIM2 were observed (Fig. 2A-B). Further, time course studies demonstrated that PIM mRNA upregulation occurs as early as 8hrs (Supplemental Fig. 1A). Differential gene expression analysis of RNA-seq data from U937 cells identified that PIM1 was among the top 20 upregulated genes in BETi treated cells compared to DMSO control (Enrichment score = -3.298) and that a gene set previously reported to be enriched in PIM overexpressing myeloid cells was also upregulated in our BETi treated cells (Fig. 2C, Supplemental Figure 1B)^32^. We next confirmed the increased PIM levels after BETi in multiple myeloid leukemia cell lines. Four of nine cell lines demonstrated increased PIM kinase protein levels at 24 hours (Supplemental Fig. 1C). While PIM upregulation was heterogeneous, the BETi dependent increases in PIM levels correlated to increased synergy with BETi and PIMi *in vitro* (Fig. 2D and E). Since current inhibitors in clinical development are pan-BET inhibitors, including those tested here, we sought to investigate which BET proteins were most associated with PIM upregulation. We individually genetically depleted BRD2, 3 and 4 in U937 and SKM1 cells and found that only BRD4 knockdown resulted in significant upregulation of PIM1 levels (Fig 2F and G) consistent with the known expression of BRD4 in the hematopoietic compartment^8,33^. Collectively, these data suggest that BET inhibition leads to increased PIM expression in a subset of cell lines that is associated with synergy between BET and PIM inhibitors.

**Figure 2.**
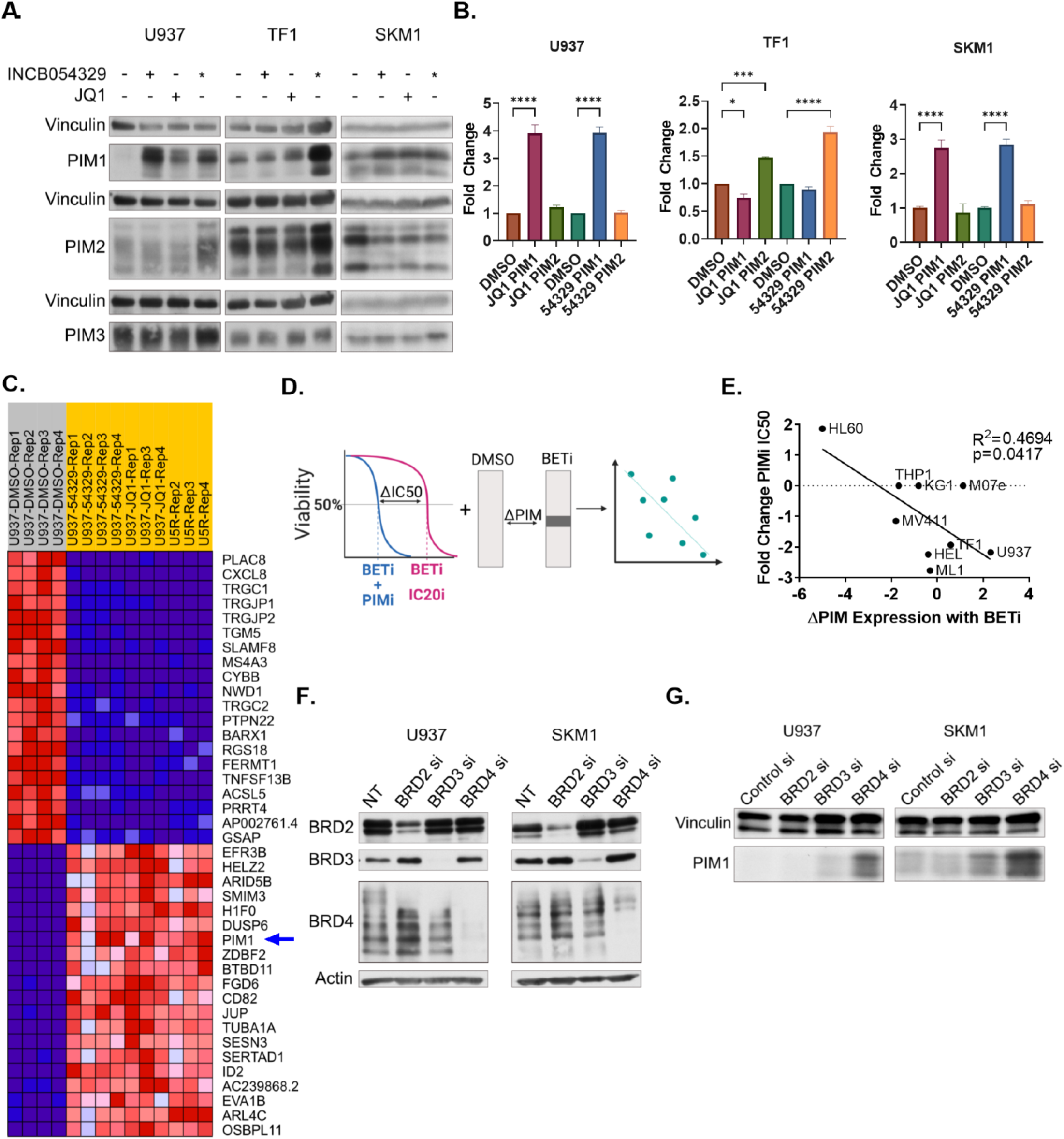
PIM kinases are increased after BET inhibition: (A) Western blot of cells lines treated with BETi for 24hrs. + indicates treatment, * indicates persistent cell lines. Each PIM kinase was run on a separate gel due to similar sizes and combined to produce the figure. (B) qPCR of cell lines treated with BETi for 24hrs. (C) GSEA analysis of BETi treated U937 cells showing the top 20 up and downregulated genes. Red = upregulated, Blue = downregulated. (D) Graphic detailing the method for generating the data in Fig. 2E. Figure created in Biorender. (E) Correlation plot of PIMi IC_50_ and PIM kinase changes of cells treated with BETi for 24hrs. (F) Western blot of BET family proteins in cells treated with siRNA against each individual BET protein. (G) Western blot of PIM1 in cells treated with siRNA against BET proteins. For both F and G, BET proteins were run on separate gels due to similar sizes and combined to produce the figure. Actin was used instead of vinculin because BRD2 and BRD3 are very similar in size. *=p<.05, ***=p<.0005 ****=p<.00005.

### PIM1 overexpression is sufficient to induce resistance to BETi and sensitivity to PIMi

We next sought to determine whether increases in PIM1 alone could drive persistence to BET inhibition as well as contribute to the observed synergy seen *in vitro*. To test this, single cell PIM overexpressing SKM1 clones were derived by transducing a GFP expressing lentiviral vector encoding PIM1. All four SKM1 clones engineered to overexpress PIM1 were both persistent to BETi, and significantly more sensitive to PIMi (Fig. 3A-B). Moreover, PIM1 levels correlated with resistance to BET inhibition (R^2^=0.9925, p=.0037), indicating that PIM1 overexpression is sufficient for BET inhibitor persistence and sensitization to PIM inhibition *in vitro* (Fig 3C). Of note, although all PIM1 overexpressing clones were more sensitive to PIM inhibition, there was no correlation between levels of PIM1 expression and PIMi sensitivity (Supplemental Fig. 2A). Additionally, we performed *in vitro* competition assay by co-culturing SKM1 cells with two isogenic PIM1 overexpressing clones in the presence BETi or vehicle control. After 5 days of treatment with BETi, there was a statistically significant increase in PIM overexpressing isogenic cells indicating that PIM1 overexpressing cells were selected in the presence of their parental counterparts (Figure 3D). To determine whether PIM1 overexpression leads to BET inhibitor persistence and PIM sensitivity *in vivo*, heterotopic SKM1 xenograft models were generated of P1-14 SKM1 PIM overexpressing clones and isogenic controls. As in the above *in vivo* experiments, flank tumors were allowed to grow until 100-150 mm^3^ and treatment was initiated for two weeks. These experiments demonstrated statistically significant decreases in tumor volume in PIM over expressing SKM1 clones after PIM inhibition compared to parental cells suggesting that PIM overexpression is sufficient for PIM inhibitor sensitivity *in vivo* (Fig. 3E).

**Figure 3.**
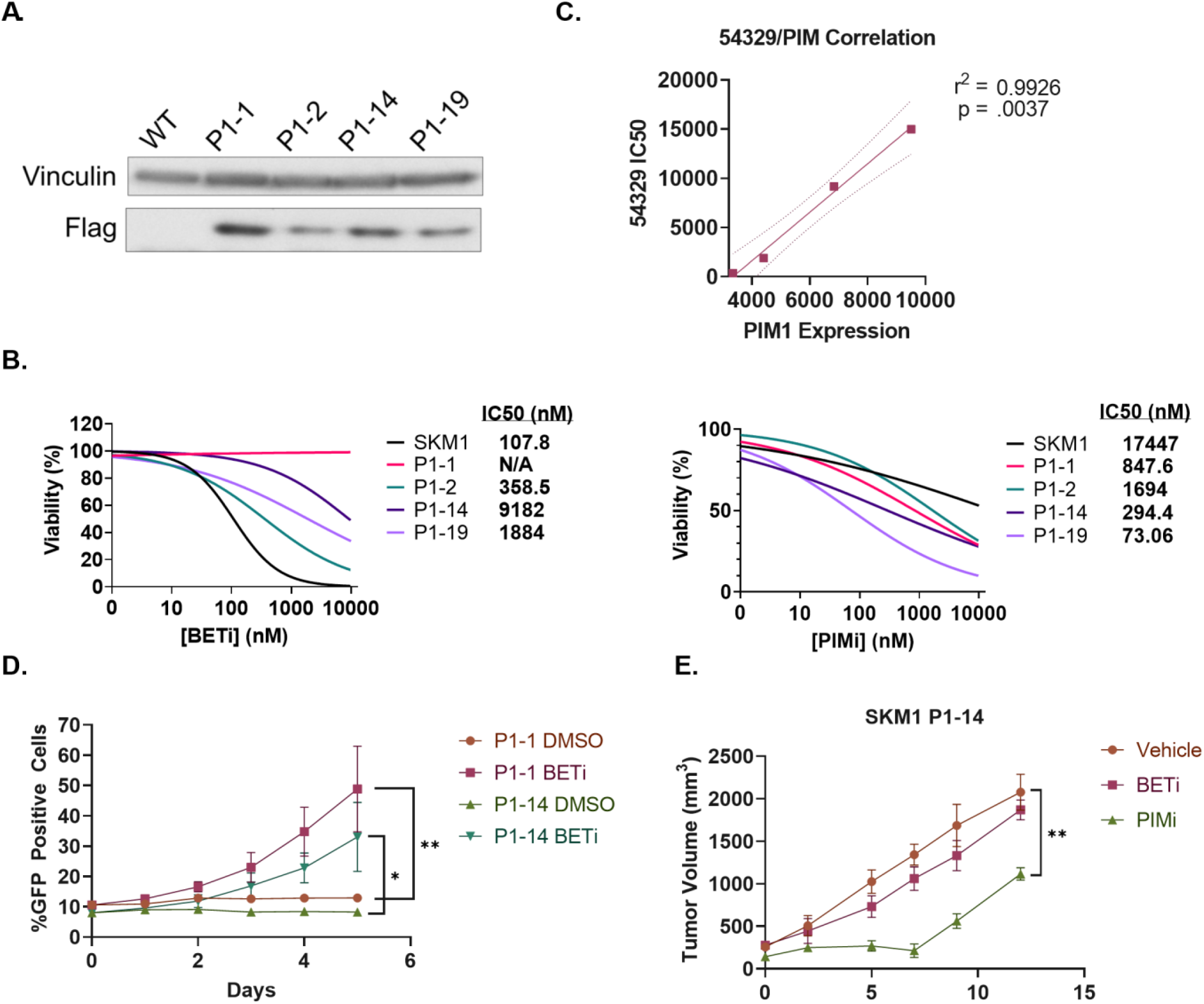
PIM1 overexpression is sufficient to induce BETi resistance and PIMi sensitivity: (A) Western blot of SKM1 cells transduced with Flag-Tagged PIM1. (B) IC_50_ curves of SKM1 cells treated with BETi (INCB054329) and PIMi(INCB053914). Cells were incubated with drug for 72hrs. (C) Correlation of PIM1 expression and BETi IC_50_ for WT and PIM1 overexpressing SKM1 cells. (D) Competition assay with SKM1 P1-1 and SKM1 P1-14 cells cultured with WT cells at a 1:10 ratio and treated with BETi for 5 days. Flow cytometry was used to determine GFP positivity. Significance was determine using AUC. (E) Tumor volumes of mice with subcutaneously implanted SKM1 P1-14 cells treated with PIMi(INCB053914). N=4 mice per group, INCB053914 N=3. Mice were euthanized when tumors reached 2cm in size. Significance was determined by comparing the AUC. *= p<.05, **=p<.005.

### BETi decreases miR-33a expression leading to increased PIM1 levels

BETi exert profound effects on the transcriptome but are generally thought to *downregulate* transcriptional activity^6,34^. Therefore, to resolve the paradoxical increase in PIM levels after treatment we first explored BET inhibitor dependent miRNA depletion hypothesizing that depletion of miRNAs that target PIM may lead to the observed increases in PIM levels. BET inhibitors can augment miRNAs via inhibition of miRNA biogenesis at super enhancer regions and/or via direct transcriptional repression of precursor RNA species^35,36^. We treated both U937 and SKM1 cells with either an Argonaute RISC Catalytic Component 2 (AGO2) inhibitor (Acriflavin) or a Dicer inhibitor (Poly-l-lysine), two central components of miRNA biogenesis, and measured protein PIM1 levels. Indeed, treatment with either AGO or Dicer inhibitors was sufficient to increase PIM1 levels across both cell lines suggesting that inhibition of miRNA biogenesis can recapitulate BET inhibitor induced PIM1 upregulation (Fig. 4A). To narrow putative miRNAs that may be responsible for PIM1 upregulation, we used the computational approach outlined in Figure 4B. Briefly, miRNAs were identified by cross referencing putative PIM1 binding miRNA from the microRNA Data Integration Portal (miRDIP), miRNA with super enhancers from Suzuki et al. and published PIM1 interacting miRNA^37-41^. This led to the identification of 4 putative miRNAs whose expression was evaluated after BET inhibitor treatment. Of these, miR-33a was the only miRNA with a significant time dependent decrease after treatment with two BETi (Fig. 4C). This was consistent with whole transcriptome RNA-sequencing performed in U937 cells that demonstrated an enrichment of miR-33a targets in BETi treated cells compared to control (Fig. 4D, Enrichment Score = -0.447, FDR q=.0059). To determine if miR-33a depletion was necessary for BET dependent PIM upregulation, we electroporated a miR-33a mimic into both U937 and SKM1 cells treated with either BETi or DMSO for 24hrs and collected pellets for both RNA and protein after 48hrs (Fig. 4E). Evaluation of PIM1 protein levels demonstrated that cells with miR-33a overexpression were protected from BETi dependent PIM upregulation (Fig. 4F). These data suggest that reduced levels of miR-33a after BET inhibition leads to an increase in PIM1 expression.

**Figure 4.**
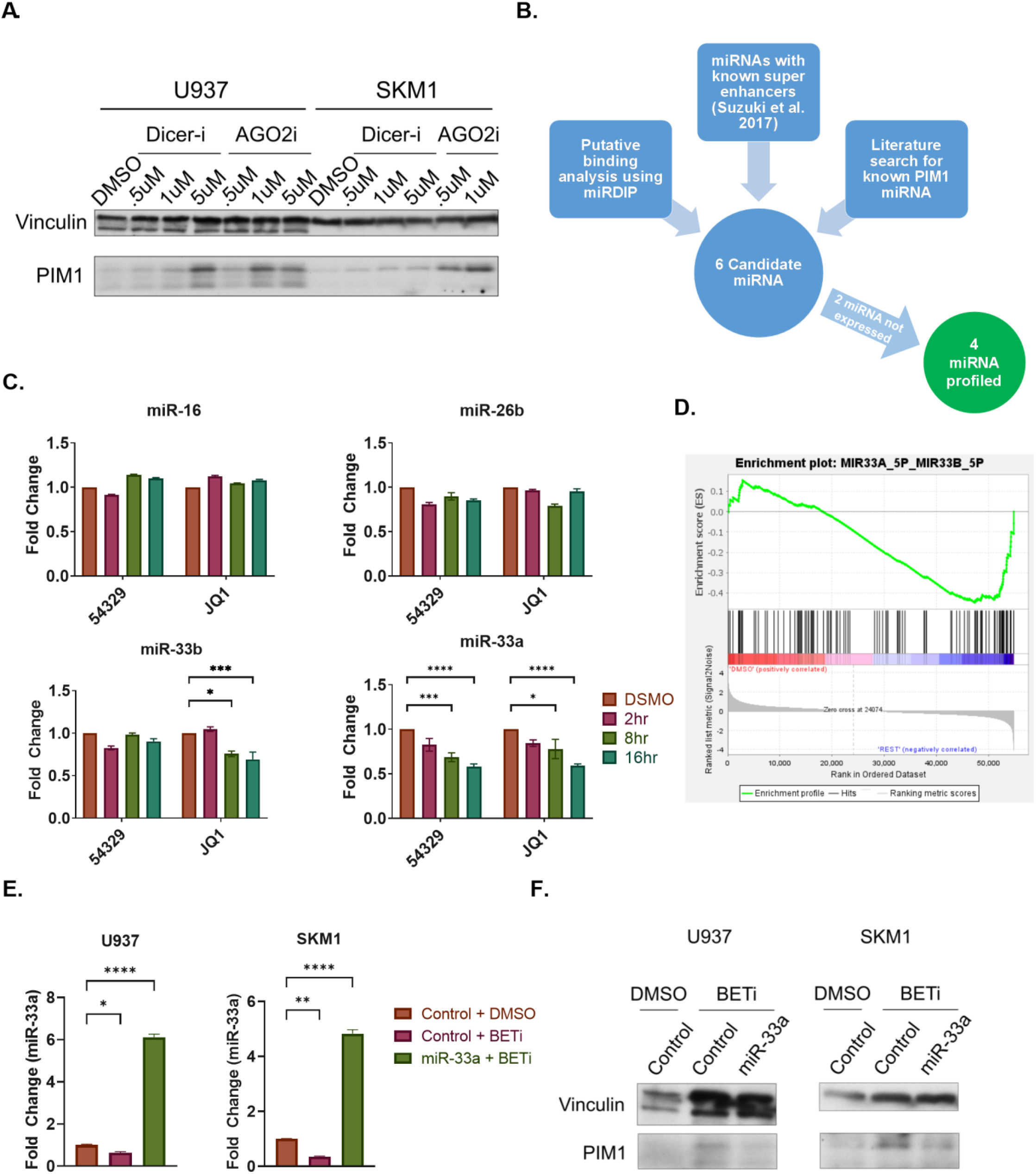
miR-33a is downregulated after BET inhibition and is responsible for PIM1 upregulation: (A) Western blots of PIM1 in cells treated with AGO2 or Dicer inhibitors. (B) Flow chart of process for selecting miRNAs for further analysis. (C) qPCR of 4 candidate miRNAs in SKM1 cells treated with BETi(INCB0543239) for 2-16hrs. (D)GSEA enrichment plot for miR-33a/miR-33b targets in U937 cells treated with BETi(INCB054329) for 24hrs. Phenotype 1 = DMSO and phenotype 2 = BETi treated/persistent cells. (E) qPCR of cells treated with both miR-33a mimic and BETi(INCB054329). (F) Western blot of cells treated with miRNA mimic and BETi(INCB054329). *= p<.05, **=p<.005, ***=p<.0005, ****=p<.00005.

Last, to explore whether BETi directly and specifically impact miR-33a we profiled transcript levels of *SREBF2* after BET inhibition as miR-33a is intronically located between exons 19 and 20 of this gene (Fig. 5A). This analysis demonstrated no difference in *SREBF2* transcript expression after treatment suggesting that BET inhibitors do not directly impact primary miR-33a transcription in leukemia cells (Fig. 5B). This was observed both with primers probing the intronic region between exons 19 and 20 as well as primers measuring total *SREBF2* (Fig. 5B). Importantly, no other promoters were identified near *SREBF2* that would transcribe miRNA-33a independently using publicly available data from CHIP-atlas (Supplemental Fig. 3)^42,43^. While the rapid turnover of miRNA precursor species precludes precise measurements of their relative abundances after treatment, we attempted to profile the range of miR-33a precursors after BET inhibitor treatment at different time points. Indeed, mature miRNA isoforms (i.e. 3p and 5p) were consistently depleted upon BET inhibitor treatment, but pre-miR-33a did not significantly decrease congruent with the postulated role of BET inhibitor repression of miRNA biogenesis^36^(Fig. 5C and D). Moreover, broad miRNA expression profiling in U937 cells demonstrated a global statistical down regulation of miRNAs in 2 replicates suggesting that miR-33a downregulation may occur through impairment of miRNA biogenesis (Figure 5E).

**Figure 5.**
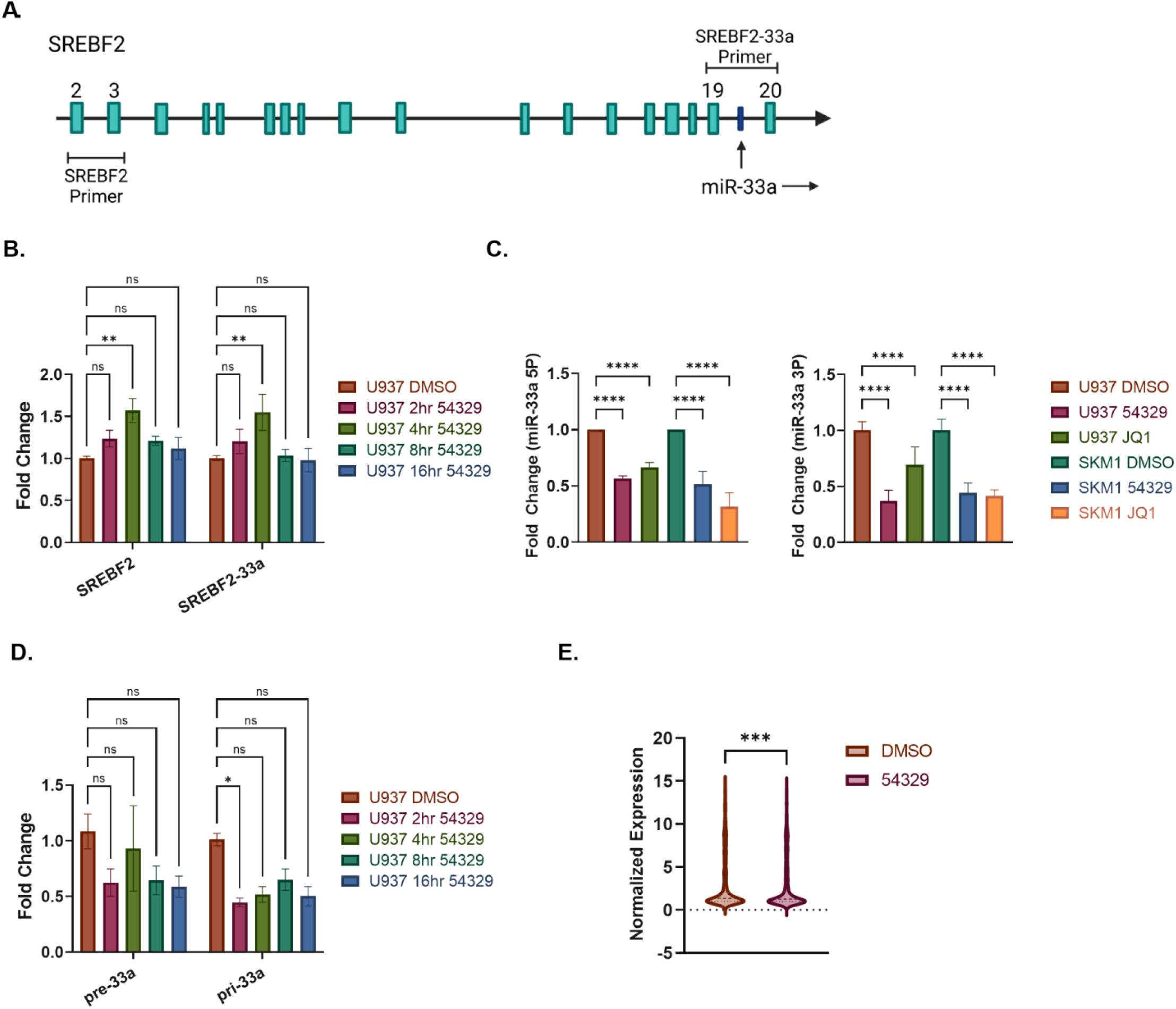
BETi inhibition of miRNA biogenesis results in miR-33a downregulation: (A) Schematic representation of miR-33a location within *SREBF2* and location of primers used in B. Figure created in Biorender. (B) qPCR of *SREBF2* levels in U937 cells treated with BETi for 2-16hrs. (C) qPCR of miR-33a-5p (Left) and miR-33a-3p (Right) levels in U937 and SKM1 cells treated with BETi for 24hrs. (D) qPCR of pre-miR-33a and pri-miR-33a levels in U937 cells treated with BETi from 2-16hrs. (E) Levels of miRNA expression in U937 cells treated with INBC054329 for 24hrs obtained from the Affymetrix GeneChip miRNA Array 4.0. *= p<.05, **=p<.005, ****=p<.00005.

### Upregulation of the GM-CSF/STAT5 axis is associated with sensitivity to combination therapy

Given the above mechanism of synergy, we hypothesized that the subset of leukemic cells which upregulated PIM after BET inhibition could be identified *a priori* by their respective pre-treatment PIM transcriptional activity. We posited that leukemia cells in a transcriptionally active state at the PIM1 loci would be primed to upregulate PIM upon BETi dependent downregulation of its inhibitory miRNAs. To explore this possibility, we measured pSTAT5 in the presence or absence of granulocyte macrophage colony stimulating factor (GM-CSF) as the GM-CSF/STAT5 axis is known to be the canonical upstream signal required for PIM transcription^44-47^. Consistent with our hypothesis, cells that exhibited BET dependent PIM upregulation demonstrated pSTAT5 activation after only 0.1ng/mL of GM-CSF stimulation (Fig. 6A). This pSTAT5 enrichment was accompanied by STAT5 occupation at the PIM1 downstream enhancer in human leukemia cells sensitive to our proposed combination therapy (Fig. 6B). GM-CSF stimulation also led to enrichment of RNA PolII at both the PIM1 enhancer and promoter in PIM upregulating cell lines but not in those leukemia cells that did not upregulate PIM (Fig. 6C). Collectively our data suggests that GM-CSF sensitive myeloid malignancies are transcriptionally primed at the PIM1 loci and associated with sensitivity to BET and PIM inhibition.

**Figure 6.**
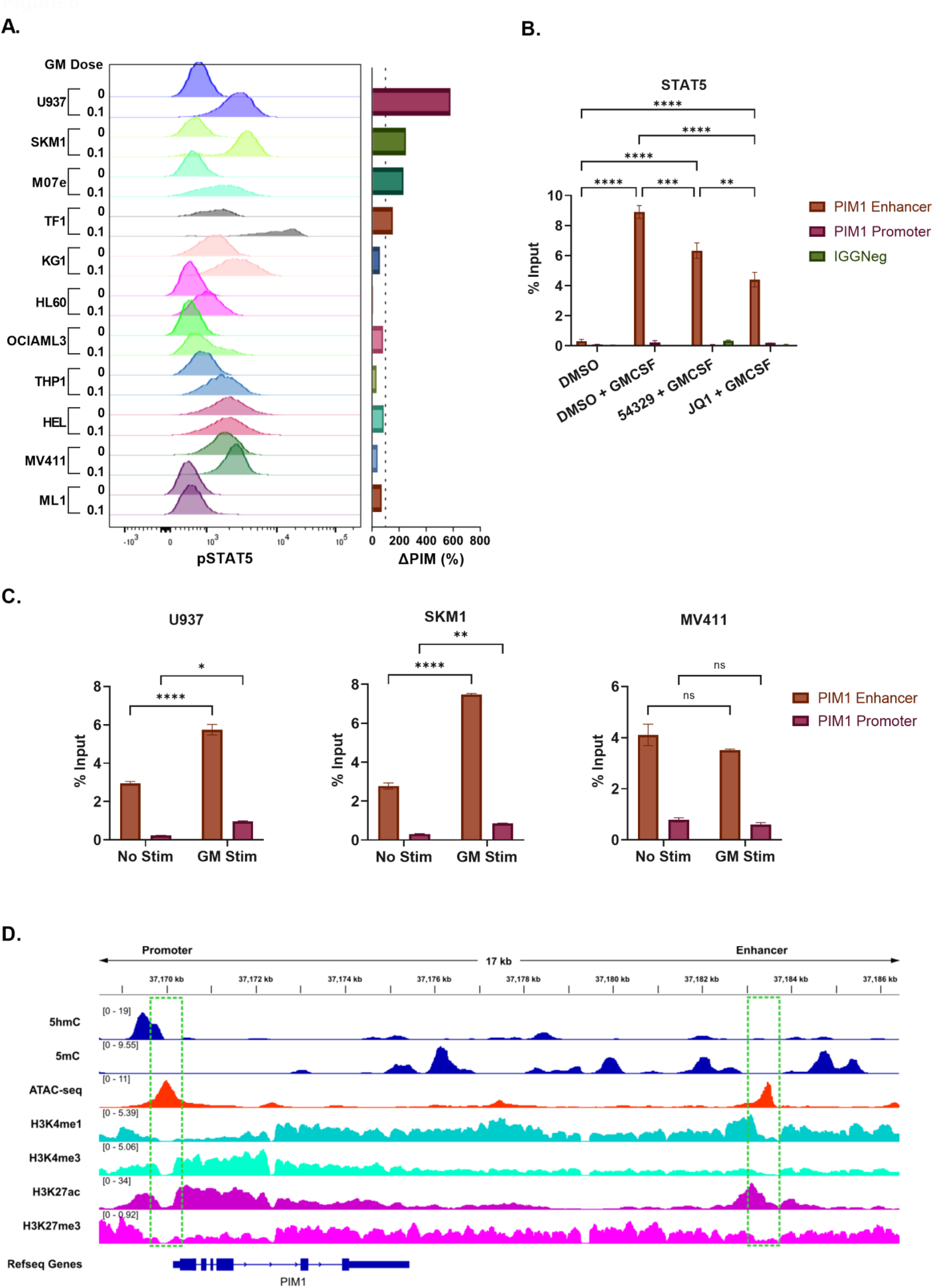
Upregulation of the GM-CSF/STAT5 axis is associated with sensitivity to combination therapy. (A) Flow cytometry analysis of pSTAT5 levels after stimulation with 0.1ng/mL GM-CSF in 11 myeloid cells lines with corresponding PIM levels after treatment with a BETi. (B) ChIP-PCR of STAT5 levels in U937 cells at the PIM1 promoter and enhancer after stimulation with GM-CSF(10ng/mL) and treatment with BETi (500nM). (C) ChIP-PCR of RNA PolII at the PIM1 promoter and enhancer in U937, SKM1 and MV411 cells after stimulation with 10ng/mL GM-CSF. (D) ChIP-seq data from 16 unique CMML patients at the PIM1 locus.

Chronic Myelomonocytic Leukemia (CMML) is a rare hematologic malignancy classified as a Myelodysplastic/Myeloproliferative overlap syndrome by the World Health Organization^48^. Clinically and pathologically, this disease is characterized by bone marrow dysplasia, peripheral monocytosis, cytopenias, and a propensity for transformation to Acute Myeloid Leukemia, all of which contribute to a poor overall survival^48^. Molecularly, CMML is hallmarked by GM-CSF hypersensitivity in a mutational and subtype independent manner ^49,50^. To determine whether this molecular feature was associated with transcriptionally primed PIM, we leveraged our previously published multi-omic epigenetic dataset of 16 CMML patients that enabled us to probe chromatin accessibility and histone marks at the PIM1 loci ^51^. Both when viewed in aggregate (Fig. 6D) or as individual patients (Supplemental Fig. 4) the PIM1 promoter and enhancer demonstrated epigenetic marks consistent with transcriptional activity supporting the notion that CMML may represent a subtype of leukemia enriched for sensitivity to BET and PIM inhibition.

### Combination therapy with BET and PIM inhibition is a viable therapeutic strategy in primary CMML cells *in vitro and in vivo*

To determine if the synergy between BET and PIM inhibitors is evident in primary CMML samples we performed clonogenicity assays with bone marrow mononuclear cells (BMMCs) from 10 unique CMML patients (Supplemental Table 3), in duplicate, treated with 100nM INCB054329 and 500nM INCB053914 or the combination. While single agent INCB054329 and INCB053914 demonstrated modest reductions of clonogenicity, a significant reduction in clonogenicity was seen with combination therapy compared to all groups. (Fig. 7A and B). To determine whether combination therapy was a viable therapeutic approach *in vivo*, we generated CMML patient derived xenografts (PDX) as previously described (Supplemental Table 3) ^52^. After engraftment was established in each model, mice were randomized (3-5 mice per group) and treated with BET inhibitor, PIM inhibitor, or the combination for 2 weeks using the same doses as heterotopic cell line xenograft experiments (Fig. 7C). Initially, mice treated with the maximum tolerated dose of BETi and PIMi rapidly lost weight and had unacceptable toxicity (data not shown). However, given that *in vitro* synergy optimally occurred at lower doses of both inhibitors and nanomolar levels of BETi were sufficient to induce PIM upregulation (Supplemental Fig. 5), PDX experiments were repeated with low dose BET and PIM inhibition. Mice treated with either low-dose INCB057643 or INCB053914 alone showed a variable response to treatment, similar to *in vitro* experiments, while the combination was consistently able to reduce leukemic engraftment as evidenced by a reduced percentage of human CD45+ cells in the bone marrow ^53-55^ by both flow (Fig. 7D) (BETi vs Combo mean rank diff. = 13.49, p=0.0049. PIMi vs Combo mean rank diff. = 11.30, p=.030) and IHC (Fig. 7E and F). Finally, we profiled PIM expression in our PDX models to determine whether the postulated mechanism of synergy occurred in primary patient samples. Immunohistochemistry (IHC) was performed on spleen sections using rabbit anti-PIM1 and anti-PIM2. To quantitate human specific PIM expression, we computationally overlaid PIM IHC with that of human CD45 (see methods)(Figure 7G). This analysis demonstrated that PIM upregulation occurred after BET inhibitor treatment *in vivo* in primary samples.

**Figure 7.**
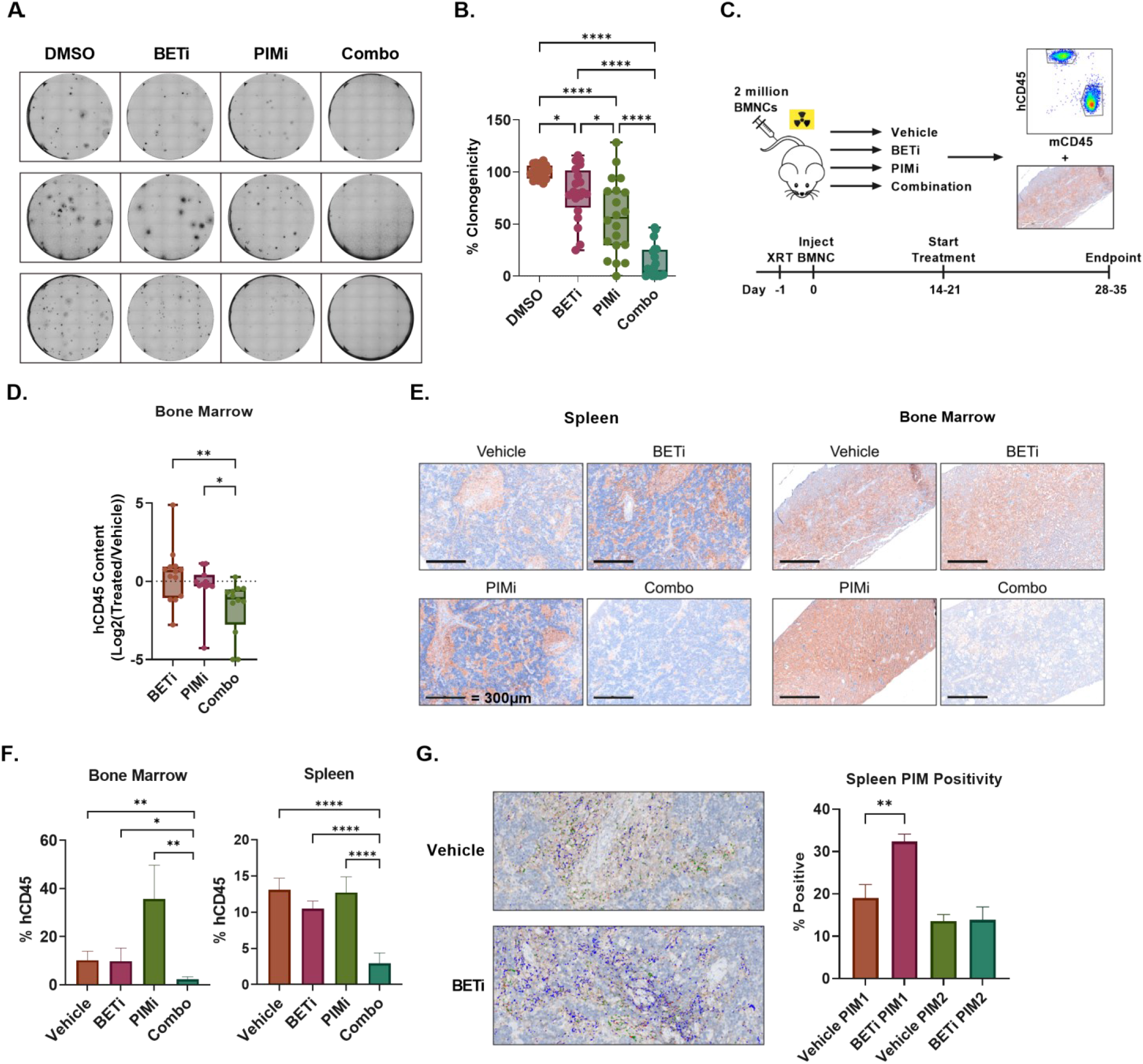
PDX models of CMML recapitulate *in vitro data*. (A) Representative images from 3 patient sample CFAs. (B) Quantification of CFA data, n=10 unique patients. (C) Graphical representation of PDX experiment timeline. (D) Flow cytometry analysis of hCD45 content in bone marrow of mice from 4 PDX experiments with 4 unique patients. Mice were treated with either BETi(INCB057643), PIMi(INCB053914) or combination. Significance determined using Kruskal-Wallis. (E) Representative images of bone marrow and spleen slides stained with hCD45. (F) Quantification of hCD45 in bone marrow and spleen IHC slides from PDX experiments. (G) Left: Representative image of a PDX spleen stained with hCD45, PIM1 and PIM2. Slides were stained with individual markers and overlaid using a computational program described in methods. Blue color represents area of hCD45 and PIM1 colocalization. Right: Quantification of the colocalization of hCD45/PIM1 and hCD45/PIM2 in spleen samples taken from PDX experiments. *=p<.05, **=p<.005, ****=p<.00005.

## Discussion

Despite advances in the molecular pathobiology and genetics of myeloid malignancies, no targeted therapeutics have demonstrated an impact on overall survival or augment natural history. This is especially evident in CMML where there are no CMML-specific approved therapies and the vast majority of patients will succumb to disease within 5 years ^56^. To address this therapeutic gap, we utilized a targeted chemical screen and identified BETi and PIMi as a synergistic combination in preclinical models *in vitro* and *in vivo*. While this synergy has not been previously reported, it is consistent with recent studies suggesting that PIM kinase upregulation may be associated with disease progression and resistance to cytotoxic therapies in AML^57-59^. It is also consistent with the notion that BET inhibitor and kinase inhibitor combination therapy may be an attractive therapeutic strategy in hematologic malignancies^60-64^.

Our study identified that PIM protein and RNA levels were paradoxically upregulated after BET inhibitor treatment in multiple cell lines and that this upregulation was necessary for sensitivity to PIM inhibition. That PIM kinase upregulation was sufficient to induce this phenotype, without upstream activation, is consistent with its known mechanism of phosphorylation. Unlike many serine threonine kinases which require a secondary phosphorylation event in order to become active, PIM kinases are constitutively active after translation^65^. Given the profound downregulation of transcriptional activity and the paradoxical increase in PIM kinase RNA in our leukemia models, we hypothesized BET inhibitors may downregulate a post-transcriptional repressor of PIM. Indeed miR-33a, a known regulator of PIM kinase, was down regulated and necessary for the observed BETi dependent PIM upregulation^37,66,67^. Further, our data strongly suggests that BETi dependent impairments in miRNA biogenesis, and not direct transcriptional repression of mir-33a precursors, underlies the mechanism of miR-33a downregulation making this proposed combination therapy mechanistically novel.

Last, we demonstrated that GM-CSF sensitive cells were associated with sensitivity to combination therapy and therefore utilized our CMML PDX models, which recapitulate many features of the human condition ^52^, to credential this therapy in a randomized murine clinical trial and identified statistically superior repression of human leukemia engraftment with combination therapy across all models. While our data provides strong evidence to clinically test low dose BET and PIM inhibition in CMML, we anticipate future clinical studies to identify the biologically effective dose of BET inhibitor that upregulates PIM kinase in humans so that responses are enriched and toxicity minimized.

## Acknowledgements

This study was supported (in part) by research funding from Incyte to C.L. and E.P., and the National Institutes of Health, National Cancer Institute (R37CA234021) and by the Flow Cytometry, Microscopy, Histology, Genomics cores, and Vivarium at the H. Lee Moffitt Cancer and Research Institute, a comprehensive cancer center designated by the National Cancer Institute (P30-CA076292).

## Authorship

Contribution: F.K. performed the compound screen under the supervision of U.R.; J.M.G and M.B. performed bioinformatics analysis of sequencing data; M.E.B., H.N., A.V. and T.K. assisted with in vivo work; M.M.P., M.B., T.L.L, C.M.F. and A.G.M. contributed and analyzed ChIP-seq data. C.L. and E.P. designed the experiments, analyzed the results and wrote the manuscript. All authors revised and approved the final manuscript.

## Materials and Methods

### Experimental Models and Subject Details

#### Cell Lines

U937, MV4-11, SKM-1, OCI-AML-3, HEL, HL-60, THP1, ML1 and KG1 cells were cultured in RPMI with 10% fetal bovine serum (FBS). TF1 Cells were cultured in RPMI with 10% FBS and 2ng/mL GM-CSF. M07-e cells were cultured in RPMI with 10% FBS and 10 ng/mL GM-CSF.

#### Heterotopic Cell Line Models and CMML PDX

All animal studies were approved by the Moffitt Cancer Center Institutional Animal Care and Use Committee.

U937, SKM1 or SKM1 P1-14 cells were resuspended in cold 0.9% sterile saline and mixed with Matrigel Matrix to a final protein concentration of 7mg/mL. 3×10^5^ U937 or 1×10^6^ SKM1 and SKM1 P1-14 cells were injected into the right flank of NGS-S mice and allowed to reach between 100mm^3^ and 150mm^3^ before beginning treatment. Tumors were measured at least twice a week by caliper and tumor volume was calculated using the formula; Tumor volume = width × width × length × 0.52. Mice were divided into 4 groups: vehicle, INCB057643, INCB053914 or Combination. INCB057643 was administered once a day at 10mg/kg, 5 days a week by oral gavage. INCB053914 was administered twice a day at 30mg/kg, 5 days a week by oral gavage. Both compounds were dissolved in 5% N,N-dimethylacetimide/95% 0.5% methylcellulose.

For CMML PDX experiments, frozen BMMCs were first thawed and treated with DNAse I for 15 minutes to create a single cell suspension. Cells were washed once and resuspended in 0.9% sterile saline and injected via tail vein into NSG-S mice sub-lethally irradiated the day prior. At least 2 million cells were injected into each mouse and treatment started between 2-3 weeks after injection. Mice were divided into the same groups as the heterotopic cell line models. Treatment lasted 2 weeks, after which mice were euthanized and the spleen, peripheral blood and femur taken for analysis. One femur and a portion of the spleen were fixed in formalin and used for IHC. Another portion of the spleen, peripheral blood and bone marrow were further processed by creating a single cell suspension and lysing red blood cells with ACK lysis buffer. Cells were then washed with PBS and stained with zombie violet viability dye (Biolegend) before fixation in 1.6% formaldehyde and storage at 4°C.

### Methods

#### Viability Assays

For the drug screen, cells were plated with compounds in 384 well plates and viability was assessed after 72hrs using Cell-Titre Glo (Promega) according to the manufacturer’s instructions. For all other viability assays, cells were plated with compounds in 96 well plates and viability was assessed after 72hrs using CCK8 following the manufacturer’s instructions. synergy was calculated using Zero Interaction Potential (ZIP) via SynergyFinder^68^

#### Persistent Cell Lines

U937 and TF-1 cells were grown in medium containing 500nM INCB054329 and SKM-1 cells were grown in medium containing 300nM INCB054329. Persistence was determined by significantly increased IC_50_ by CCK8 and steady growth in medium containing INCB054329.

#### Colony Forming Assays

Frozen BMMCs were thawed and prepared in a similar manner to PDX experiments. Cells were then resuspended in IMDM + 2% FBS at a concentration of 200,000 cells per mL. 300µL of cell suspension and 3μL of each compound were added to 3mL Methocult 4034 from StemCell and mixed by vortexing for 1 minute. 1.1 mL of cell mixture was plated in StemCell smart dishes, incubated for 14 days and read on StemVision (Stem Cell Technologies) for the final colony count.

#### Western Blotting

All cells were lysed using RIPA lysis buffer and protein quantified using BCA. SDS– polyacrylamide gel electrophoresis was performed using 7.5, 10 or 12.5% bis-tris gels and protein was transferred to PVDF membranes using a wet transfer system (Bio-Rad). Membranes were blocked with 5% milk in TBS-T and incubated overnight with primary antibody in either milk or BSA at manufacturer recommended concentrations. Blots were washed multiple times in TBS-T before addition of HRP-conjugated secondary antibody diluted in 5% milk and incubated for 1hr at room temperature. Antibodies used: BRD2, BRD4, FLAG-TAG, PIM1, PIM2, PIM3(Cell Signaling), BRD3(Bethyl), Actin(Sigma-Aldrich), Vinculin(Sigma-Aldrich).

#### RNA Extraction

Total RNA from cultured cells was extracted using either Quick-RNA Miniprep (Zymo Research) for gene expression or miRNeasy/miRNeasy advanced (Qiagen) for miRNA detection.

#### q-RT-PCR

RNA was converted into cDNA using iScript® Reverse Transcription Supermix for RTqPCR (Bio-Rad). qRT-PCR reactions were performed in triplicate using probes designed and ordered from IDT, or off the shelf TaqMan assays (Applied Biosystems). For miRNA, cDNA was generated using the TaqMan Advanced miRNA cDNA Synthesis Kit. qRT-PCR reactions were also performed in triplicate using off the shelf TaqMan Advanced miRNA Assays (Applied Biosystems).

#### ChIP-PCR

U937, SKM1, or MV411 cells were serum starved overnight. The next day, cells were stimulated with 10ng/mL GM-CSF for 15mins and immediately fixed with 1% formaldehyde for 10mins. Formaldehyde was quenched with glycine and cells were washed 2X with cold PBS before being snap frozen on dry ice and stored at -80ºC. Fixed cells were then prepared using the SimpleChIP Magnetic Bead Kit (Cell Signaling Technologies) according to manufacturer’s recommendations. DNA was sheared using a Qsonica Q800R3 with the following settings: 50% amplitude, 30sec pulse, 5min shearing time. DNA was sheared 5min, spun down, and sheared an additional 5min. Antibodies for STAT5 (Cell signaling technologies), and RNA PolII(Active Motif) were incubated overnight before continuing with the protocol according to manufacturer’s recommendations.

#### RNA-seq and GSEA

U937 cells were treated with DMSO, JQ1 and INCB054329 for 24hrs in quadruplicate. Persistent cells treated with INCB054329 were also included in quadruplicate. RNA was extracted and screened for quality on an Agilent BioAnalyzer. The samples were then processed for RNA-sequencing using the Nugen Universal RNA-seq kit(NuGEN). Briefly, 100 ng of RNA was used to generate cDNA and a strand-specific library following the manufacturer’s protocol. Quality control steps including BioAnalyzer library assessment and quantitative RT-PCR for library quantification were performed. Two libraries failed QC and were excluded. The libraries were then sequenced the Illumina NextSeq 500 v2 sequencer with two high-output 75-base paired-end runs in order to generate approximately 25 to 30 million read pairs per sample. Sequencing data was mapped to hg38 using STAR “Spliced Transcripts Alignment to a Reference”^69^. Raw data was cleaned by removing any genes with less than 10 reads or present in less than half of the samples before running differential analysis through DESeq2^70^. Normalized counts were run through GSEA 4.1.0 with default parameters except permutation type, which was set to gene_set^71^.

#### Transduction of Cells with PIM1

SKM1 cells were transduced with a Flag-Tagged, 34kDa isoform of PIM1 in a pCDH-CMV-MCS-EF1α-GreenPuro Cloning and Expression Lentivector(System Biosciences) via the Spinfection method. Briefly, cells were resuspended in Opti-MEM and plated into 6-well plates along with fresh virus, Lipofectamine-2000 and polybrene. Cells were centrifuged for 90mins at 2200rpm in a 37°C centrifuge, and incubated at 37°C for 1 hour, after which 1 mL of normal growth medium was added and cells were incubated overnight. Cells were then centrifuged and resuspended in normal growth medium. After 1 week, cells were single cell sorted for GFP positivity. Single cell clones were profiled for successful transduction by western blotting for Flag-tag.

#### Electroporation

The Neon Transfection System(ThermoFisher Scientific) with 100μL tips was used to deliver siRNA or miRNA mimics. Cells were first washed with PBS and resuspended in R buffer at a concentration of 5×10^7^ cells per mL. siRNA or miRNA mimics were added to a final concentration of 5μM, mixed thoroughly and cells were electroporated with the following settings: 1400V, 10 pulse width, 3 pulses. Cells were then added to 10mL RPMI with 10% FBS and incubated for either 48 (miRNA mimics) or 72 (siRNA) hours before collection for qPCR and western blotting. For experiments with miRNA mimics, INCB054329 was added 24hrs after electroporation.

#### miRNA Array

RNA was extracted from U937 cells treated with DMSO, INBC054329 or JQ1 for 24hrs using miRNeasy Advanced kit (Qiagen). Thermo GeneChip miRNA 4.0 arrays were processed and hybridized according to the manufacturer’s protocol (ThermoFisher Scientific, Waltham, MA). Briefly, 500ng of RNA was processed using the FlashTag Biotin HSR RNA Labeling Kit and following poly-adenylation and ligation of a biotinylated RNA tag, the product was hybridized to GeneChip miRNA 4.0 arrays at 48C for 16 hours at 60 RPM using the GeneChip Hybridization Oven 645. The hybridized miRNA arrays were then washed and stained using the GeneChip Fluidics Station 450, followed by scanning on the Thermo GeneChip Scanner 3000 7G. Data were reviewed for quality control and analysis was performed using the GeneChip Transcriptome Analysis Console v4.0.

#### Competition Assays

SKM1 P1-1 or P1-14 were plated with SKM1 WT cells in a 1:10 ratio. The initial mixture of cells was checked before any treatment started. Cells were plated with 150nM INCB054329 and incubated for 5 days. Each day, 1mL of cell suspension was taken out for analysis of GFP positive cells and replaced with fresh medium with INCB054329 or DMSO.

#### Flow Cytometry

For SKM1 competition assays, live cells were washed with FACS buffer and stained with DAPI before running on a FACSCanto (BD Biosciences). For GM-CSF stimulation experiments, cells were starved overnight, incubated with zombie violet for 20mins, washed, and stimulated with varying concentrations of GM-CSF for 15 minutes. Immediately after stimulation, paraformaldehyde was added to a final concentration of 1.6% and cells were fixed for 10 mins. Cells were then washed and permeabilized using 2mL of ice-cold 95% methanol. After washing off methanol, cells were stored in FACS buffer until analysis. On the day of analysis, cells were stained with pSTAT5 antibody (BD Biosciences) for 15 minutes, washed and run on a FACSCanto. For PDX experiments, cells were resuspended in 50μL FACS buffer with 2μL of both human and mouse FCR blocking antibody and incubated for 10mins. An antibody cocktail comprised of hCD45, mCD45, hCD3, hCD33, and hCD34(BD Biosciences) was added to each tube, incubated for 15 minutes and washed with FACS buffer. Cells were run on an LSRII (BD Biosciences). Data was analyzed in FlowJo.

#### Immunohistochemistry

Slides were stained using a Ventana Discovery XT automated system (Ventana Medical Systems). Briefly, slides were deparaffinized with Discovery Wash solution and heat-induced antigen retrieval method was used in Ribo CC. Rabbit primary antibodies for hCD45(#ab10558, Abcam), PIM1(#PA5-22315, Invitrogen) and PIM2 (#710504, Invitrogen) were used in Dako antibody diluent (Carpenteria) and incubated for 60, 32 and 32 min respectively. Slides were then stained with anti-rabbit secondary (Ventana). Detection was performed using the Ventana ChromoMap kit and slides were counterstained Hematoxylin. Slides were then dehydrated and coverslipped as per normal laboratory protocol.

#### Microscopy Analysis

Serial slide sections stained for CD45, PIM1, and PIM2 we scanned with a Leica Aperio AT2 digital Pathology Slide Scanner (Leica Biosystems, Vista, CA) with a 20x/0.7NA objective lens. SVS image files were imported into Visiopharm version 2022.02 (Visiopharm A/S, Denmark) where the Tissuealign tool was used to co-register images for the 3 IHC biomarkers. After alignment, the software’s manual drawing tool was used to select Regions of Interest (ROIs) on each aligned image set and a simple intensity threshold segmentation was applied to the ROIs in order to label each co-registered image pixel into staining categories. All thresholds and settings for pixel labeling were identical for each image analyzed within each experiment.

#### Statistical Analysis

Statistical analyses and graphical presentations were performed using Prism 9.0 (GraphPad). One or Two-way ANOVA is used for calculating significance unless otherwise specified. Power analyses were used to determine number of mice in *in vivo* experiments.

#### Data Sharing Statement

For original data, please contact.

## Supplemental Figure Legends

**Supplemental Figure S1:** (A) qPCR data for *PIM1* in U937 cells treated with BETi from 2-16hr. (B) GSEA enrichment plot of RNA-seq data generated from U937 cells treated with BETi for 24hrs. (C) Western blot of PIM1, 2 and 3 in cell lines associated with the correlation plot in Figure 2E. Each PIM kinase was run on a separate gel and combined to produce the figure.

**Supplemental Figure S2:** (A) Correlation plot for PIM1 overexpressing SKM1 cells treated with PIMi.

**Supplemental Figure S3:** H3K27ac and ATAC-seq binding probabilities along *SREBF2*. Data was acquired from the ChIP-Atlas web portal and imported into IGV to generate the figure.

**Supplemental Figure S4:** Individual patient sample ChIP-Seq tracks.

**Supplemental Figure S5:** Western blot images of U937 and SKM1 cells treated with increasing does of INCB054329.

**Supplemental Table 1: Table of Compounds Used in Initial Screen**.

**Supplemental Table 2: Compounds and associated references used in filtering**.

**Supplemental Table 3: Patient characteristics for CFA and PDX experiments**.

## References

1. Patnaik, M.M., and Lasho, T. (2020). Evidence-Based Minireview: Myelodysplastic syndrome/myeloproliferative neoplasm overlap syndromes: a focused review. Hematology 2020, 460–464. 10.1182/hematology.2020000163.

2. Mason, C.C., Khorashad, J.S., Tantravahi, S.K., Kelley, T.W., Zabriskie, M.S., Yan, D., Pomicter, A.D., Reynolds, K.R., Eiring, A.M., Kronenberg, Z., et al. (2016). Age-related mutations and chronic myelomonocytic leukemia. Leukemia 30, 906–913. 10.1038/leu.2015.337.

3. Ricci, C., Fermo, E., Corti, S., Molteni, M., Faricciotti, A., Cortelezzi, A., Lambertenghi Deliliers, G., Beran, M., and Onida, F. (2010). RAS Mutations Contribute to Evolution of Chronic Myelomonocytic Leukemia to the Proliferative Variant. Clinical Cancer Research 16, 2246–2256. 10.1158/1078-0432.Ccr-09-2112.

4. You, X., Liu, F., Binder, M., Vedder, A., Lasho, T., Wen, Z., Gao, X., Flietner, E., Rajagopalan, A., Zhou, Y., et al. (2022). Asxl1 loss cooperates with oncogenic Nras in mice to reprogram the immune microenvironment and drive leukemic transformation. Blood 139, 1066–1079. 10.1182/blood.2021012519.

5. Filippakopoulos, P., Qi, J., Picaud, S., Shen, Y., Smith, W.B., Fedorov, O., Morse, E.M., Keates, T., Hickman, T.T., Felletar, I., et al. (2010). Selective inhibition of BET bromodomains. Nature 468, 1067. 10.1038/nature09504 https://www.nature.com/articles/nature09504#supplementary-information.

6. Muhar, M., Ebert, A., Neumann, T., Umkehrer, C., Jude, J., Wieshofer, C., Rescheneder, P., Lipp, J.J., Herzog, V.A., Reichholf, B., et al. (2018). SLAM-seq defines direct gene-regulatory functions of the BRD4-MYC axis. Science. 10.1126/science.aao2793.

7. Loven, J., Hoke, H.A., Lin, C.Y., Lau, A., Orlando, D.A., Vakoc, C.R., Bradner, J.E., Lee, T.I., and Young, R.A. (2013). Selective Inhibition of Tumor Oncogenes by Disruption of Super-Enhancers. Cell 153, 320–334. 10.1016/j.cell.2013.03.036.

8. Zuber, J., Shi, J., Wang, E., Rappaport, A.R., Herrmann, H., Sison, E.A., Magoon, D., Qi, J., Blatt, K., Wunderlich, M., et al. (2011). RNAi screen identifies Brd4 as a therapeutic target in acute myeloid leukaemia. Nature 478, 524–528. 10.1038/nature10334.

9. Delmore, J.E., Issa, G.C., Lemieux, M.E., Rahl, P.B., Shi, J.W., Jacobs, H.M., Kastritis, E., Gilpatrick, T., Paranal, R.M., Qi, J., et al. (2011). BET Bromodomain Inhibition as a Therapeutic Strategy to Target c-Myc. Cell 146, 903–916. 10.1016/j.cell.2011.08.017.

10. Mertz, J.A., Conery, A.R., Bryant, B.M., Sandy, P., Balasubramanian, S., Mele, D.A., Bergeron, L., and Sims, R.J. (2011). Targeting MYC dependence in cancer by inhibiting BET bromodomains. Proceedings of the National Academy of Sciences of the United States of America 108, 16669–16674. 10.1073/pnas.1108190108.

11. Bolden, J.E., Tasdemir, N., Dow, L.E., van Es, J.H., Wilkinson, J.E., Zhao, Z., Clevers, H., and Lowe, S.W. (2014). Inducible In Vivo Silencing of Brd4 Identifies Potential Toxicities of Sustained BET Protein Inhibition. Cell reports 8, 1919–1929. 10.1016/j.celrep.2014.08.025.

12. Fong, C.Y., Gilan, O., Lam, E.Y.N., Rubin, A.F., Ftouni, S., Tyler, D., Stanley, K., Sinha, D., Yeh, P., Morison, J., et al. (2015). BET inhibitor resistance emerges from leukaemia stem cells. Nature 525, 538–542. 10.1038/nature14888 http://www.nature.com/nature/journal/v525/n7570/abs/nature14888.html#supplementary-information.

13. Rathert, P., Roth, M., Neumann, T., Muerdter, F., Roe, J.-S., Muhar, M., Deswal, S., Cerny-Reiterer, S., Peter, B., Jude, J., et al. (2015). Transcriptional plasticity promotes primary and acquired resistance to BET inhibition. Nature 525, 543–547. 10.1038/nature14898 http://www.nature.com/nature/journal/v525/n7570/abs/nature14898.html#supplementary-information.

14. Wyce, A., Matteo, J.J., Foley, S.W., Felitsky, D.J., Rajapurkar, S.R., Zhang, X.-P., Musso, M.C., Korenchuk, S., Karpinich, N.O., Keenan, K.M., et al. (2018). MEK inhibitors overcome resistance to BET inhibition across a number of solid and hematologic cancers. Oncogenesis 7, 35. 10.1038/s41389-018-0043-9.

15. Shu, S., Lin, C.Y., He, H.H., Witwicki, R.M., Tabassum, D.P., Roberts, J.M., Janiszewska, M., Huh, S.J., Liang, Y., Ryan, J., et al. (2016). Response and resistance to BET bromodomain inhibitors in triple negative breast cancer. Nature 529, 413–417. 10.1038/nature16508.

16. Kurimchak, A.M., Shelton, C., Duncan, K.E., Johnson, K.J., Brown, J., O’Brien, S., Gabbasov, R., Fink, L.S., Li, Y., Lounsbury, N., et al. (2016). Resistance to BET Bromodomain Inhibitors Is Mediated by Kinome Reprogramming in Ovarian Cancer. Cell reports 16, 1273–1286. 10.1016/j.celrep.2016.06.091.

17. Kleppe, M., Koche, R., Zou, L., van Galen, P., Hill, C.E., Dong, L., De Groote, S., Papalexi, E., Hanasoge Somasundara, A.V., Cordner, K., et al. (2018). Dual Targeting of Oncogenic Activation and Inflammatory Signaling Increases Therapeutic Efficacy in Myeloproliferative Neoplasms. Cancer Cell 33, 29-43.e27. 10.1016/j.ccell.2017.11.009.

18. Faiao-Flores, F., Emmons, M.F., Durante, M.A., Kinose, F., Saha, B., Fang, B., Koomen, J.M., Chellappan, S.P., Maria-Engler, S.S., Rix, U., et al. (2019). HDAC Inhibition Enhances the In Vivo Efficacy of MEK Inhibitor Therapy in Uveal Melanoma. Clin Cancer Res 25, 5686–5701. 10.1158/1078-0432.CCR-18-3382.

19. Baker, E.K., Taylor, S., Gupte, A., Sharp, P.P., Walia, M., Walsh, N.C., Zannettino, A.C., Chalk, A.M., Burns, C.J., and Walkley, C.R. (2015). BET inhibitors induce apoptosis through a MYC independent mechanism and synergise with CDK inhibitors to kill osteosarcoma cells. Sci Rep 5, 10120. 10.1038/srep10120.

20. Sun, B., Shah, B., Fiskus, W., Qi, J., Rajapakshe, K., Coarfa, C., Li, L., Devaraj, S.G., Sharma, S., Zhang, L., et al. (2015). Synergistic activity of BET protein antagonist-based combinations in mantle cell lymphoma cells sensitive or resistant to ibrutinib. Blood 126, 1565–1574. 10.1182/blood-2015-04-639542.

21. Tomska, K., Kurilov, R., Lee, K.S., Hullein, J., Lukas, M., Sellner, L., Walther, T., Wagner, L., Oles, M., Brors, B., et al. (2018). Drug-based perturbation screen uncovers synergistic drug combinations in Burkitt lymphoma. Sci Rep 8, 12046. 10.1038/s41598-018-30509-3.

22. Cortiguera, M.G., Garcia-Gaipo, L., Wagner, S.D., Leon, J., Batlle-Lopez, A., and Delgado, M.D. (2019). Suppression of BCL6 function by HDAC inhibitor mediated acetylation and chromatin modification enhances BET inhibitor effects in B-cell lymphoma cells. Sci Rep 9, 16495. 10.1038/s41598-019-52714-4.

23. Enssle, J.C., Boedicker, C., Wanior, M., Vogler, M., Knapp, S., and Fulda, S. (2018). Co-targeting of BET proteins and HDACs as a novel approach to trigger apoptosis in rhabdomyosarcoma cells. Cancer Lett 428, 160–172. 10.1016/j.canlet.2018.04.032.

24. Mazur, P.K., Herner, A., Mello, S.S., Wirth, M., Hausmann, S., Sanchez-Rivera, F.J., Lofgren, S.M., Kuschma, T., Hahn, S.A., Vangala, D., et al. (2015). Combined inhibition of BET family proteins and histone deacetylases as a potential epigenetics-based therapy for pancreatic ductal adenocarcinoma. Nat Med 21, 1163–1171. 10.1038/nm.3952.

25. Schafer, J.M., Lehmann, B.D., Gonzalez-Ericsson, P.I., Marshall, C.B., Beeler, J.S., Redman, L.N., Jin, H., Sanchez, V., Stubbs, M.C., Scherle, P., et al. (2020). Targeting MYCN-expressing triple-negative breast cancer with BET and MEK inhibitors. Sci Transl Med 12. 10.1126/scitranslmed.aaw8275.

26. Tiago, M., Capparelli, C., Erkes, D.A., Purwin, T.J., Heilman, S.A., Berger, A.C., Davies, M.A., and Aplin, A.E. (2020). Targeting BRD/BET proteins inhibits adaptive kinome upregulation and enhances the effects of BRAF/MEK inhibitors in melanoma. Br J Cancer 122, 789–800. 10.1038/s41416-019-0724-y.

27. Derenzini, E., Mondello, P., Erazo, T., Portelinha, A., Liu, Y., Scallion, M., Asgari, Z., Philip, J., Hilden, P., Valli, D., et al. (2018). BET Inhibition-Induced GSK3beta Feedback Enhances Lymphoma Vulnerability to PI3K Inhibitors. Cell Rep 24, 2155–2166. 10.1016/j.celrep.2018.07.055.

28. Stratikopoulos, E.E., Dendy, M., Szabolcs, M., Khaykin, A.J., Lefebvre, C., Zhou, M.M., and Parsons, R. (2015). Kinase and BET Inhibitors Together Clamp Inhibition of PI3K Signaling and Overcome Resistance to Therapy. Cancer Cell 27, 837–851. 10.1016/j.ccell.2015.05.006.

29. Wunderlich, M., Chou, F.S., Link, K.A., Mizukawa, B., Perry, R.L., Carroll, M., and Mulloy, J.C. (2010). AML xenograft efficiency is significantly improved in NOD/SCID-IL2RG mice constitutively expressing human SCF, GM-CSF and IL-3. Leukemia 24, 1785–1788. 10.1038/leu.2010.158.

30. Warfel, N.A., and Kraft, A.S. (2015). PIM kinase (and Akt) biology and signaling in tumors. Pharmacology & therapeutics 151, 41–49. http://dx.doi.org/10.1016/j.pharmthera.2015.03.001.

31. Nawijn, M.C., Alendar, A., and Berns, A. (2011). For better or for worse: the role of Pim oncogenes in tumorigenesis. Nat Rev Cancer 11, 23–34.

32. Xin, G., Chen, Y., Topchyan, P., Kasmani, M.Y., Burns, R., Volberding, P.J., Wu, X., Cohn, A., Chen, Y., Lin, C.W., et al. (2021). Targeting PIM1-Mediated Metabolism in Myeloid Suppressor Cells to Treat Cancer. Cancer Immunol Res 9, 454–469. 10.1158/2326-6066.CIR-20-0433.

33. Shimamura, T., Chen, Z., Soucheray, M., Carretero, J., Kikuchi, E., Tchaicha, J.H., Gao, Y., Cheng, K.A., Cohoon, T.J., Qi, J., et al. (2013). Efficacy of BET bromodomain inhibition in Kras-mutant non-small cell lung cancer. Clinical cancer research : an official journal of the American Association for Cancer Research 19, 6183–6192. 10.1158/1078-0432.CCR-12-3904.

34. Delmore, Jake E., Issa Ghayas C., Lemieux Madeleine E., Rahl Peter B., Shi, J., Jacobs Hannah M., Kastritis, E., Gilpatrick, T., Paranal Ronald M., Qi, J., et al. (2011). BET Bromodomain Inhibition as a Therapeutic Strategy to Target c-Myc. Cell 146, 904–917. https://doi.org/10.1016/j.cell.2011.08.017.

35. Zhao, Y., Liu, Q., Acharya, P., Stengel Kristy R., Sheng, Q., Zhou, X., Kwak, H., Fischer Melissa A., Bradner James E., Strickland Stephen A., et al. (2016). High-Resolution Mapping of RNA Polymerases Identifies Mechanisms of Sensitivity and Resistance to BET Inhibitors in t(8;21) AML. Cell Reports 16, 2003–2016. 10.1016/j.celrep.2016.07.032.

36. Suzuki, H.I., Young, R.A., and Sharp, P.A. (2017). Super-Enhancer-Mediated RNA Processing Revealed by Integrative MicroRNA Network Analysis. Cell 168, 1000-1014.e1015. http://dx.doi.org/10.1016/j.cell.2017.02.015.

37. Thomas, M., Lange-Grünweller, K., Weirauch, U., Gutsch, D., Aigner, A., Grünweller, A., and Hartmann, R.K. (2011). The proto-oncogene Pim-1 is a target of miR-33a. Oncogene 31, 918. 10.1038/onc.2011.278 https://www.nature.com/articles/onc2011278#supplementary-information.

38. Liu, Y., Zhang, J., Xing, C., Wei, S., Guo, N., and Wang, Y. (2018). miR-486 inhibited osteosarcoma cells invasion and epithelial-mesenchymal transition by targeting PIM1. Cancer Biomark 23, 269–277. 10.3233/cbm-181527.

39. Deng, D., Wang, L., Chen, Y., Li, B., Xue, L., Shao, N., Wang, Q., Xia, X., Yang, Y., and Zhi, F. (2016). MicroRNA-124-3p regulates cell proliferation, invasion, apoptosis, and bioenergetics by targeting PIM1 in astrocytoma. Cancer Sci 107, 899–907. 10.1111/cas.12946.

40. Kim, K.T., Carroll, A.P., Mashkani, B., Cairns, M.J., Small, D., and Scott, R.J. (2012). MicroRNA-16 is down-regulated in mutated FLT3 expressing murine myeloid FDC-P1 cells and interacts with Pim-1. PLoS One 7, e44546. 10.1371/journal.pone.0044546.

41. Pang, W., Tian, X., Bai, F., Han, R., Wang, J., Shen, H., Zhang, X., Liu, Y., Yan, X., Jiang, F., and Xing, L. (2014). Pim-1 kinase is a target of miR-486-5p and eukaryotic translation initiation factor 4E, and plays a critical role in lung cancer. Mol Cancer 13, 240. 10.1186/1476-4598-13-240.

42. Oki, S., Ohta, T., Shioi, G., Hatanaka, H., Ogasawara, O., Okuda, Y., Kawaji, H., Nakaki, R., Sese, J., and Meno, C. (2018). ChIP-Atlas: a data-mining suite powered by full integration of public ChIP-seq data. EMBO reports 19, e46255. https://doi.org/10.15252/embr.201846255.

43. Zou, Z., Ohta, T., Miura, F., and Oki, S. (2022). ChIP-Atlas 2021 update: a data-mining suite for exploring epigenomic landscapes by fully integrating ChIP-seq, ATAC-seq and Bisulfite-seq data. Nucleic Acids Research 50, W175–W182. 10.1093/nar/gkac199.

44. Shirogane, T., Fukada, T., Muller, J.M.M., Shima, D.T., Hibi, M., and Hirano, T. (1999). Synergistic Roles for Pim-1 and c-Myc in STAT3-Mediated Cell Cycle Progression and Antiapoptosis. Immunity 11, 709–719. https://doi.org/10.1016/S1074-7613(00)80145-4.

45. Miura, O., Miura, Y., Nakamura, N., Quelle, F., Witthuhn, B., Ihle, J., and Aoki, N. (1994). Induction of tyrosine phosphorylation of Vav and expression of Pim-1 correlates with Jak2-mediated growth signaling from the erythropoietin receptor. Blood 84, 4135–4141. 10.1182/blood.V84.12.4135.bloodjournal84124135.

46. Nawijn, M.C., Alendar, A., and Berns, A. (2011). For better or for worse: the role of Pim oncogenes in tumorigenesis. Nature Reviews Cancer 11, 23–34. 10.1038/nrc2986.

47. Mikkers, H., Nawijn, M., Allen, J., Brouwers, C., Verhoeven, E., Jonkers, J., and Berns, A. (2004). Mice Deficient for All PIM Kinases Display Reduced Body Size and Impaired Responses to Hematopoietic Growth Factors. Molecular and Cellular Biology 24, 6104–6115. doi:10.1128/MCB.24.13.6104-6115.2004.

48. Arber, D.A., Orazi, A., Hasserjian, R., Thiele, J., Borowitz, M.J., Le Beau, M.M., Bloomfield, C.D., Cazzola, M., and Vardiman, J.W. (2016). The 2016 revision to the World Health Organization classification of myeloid neoplasms and acute leukemia. Blood 127, 2391–2405. 10.1182/blood-2016-03-643544.

49. Padron, E., Painter, J.S., Kunigal, S., Mailloux, A.W., McGraw, K., McDaniel, J.M., Kim, E., Bebbington, C., Baer, M., Yarranton, G., et al. (2013). GM-CSF–dependent pSTAT5 sensitivity is a feature with therapeutic potential in chronic myelomonocytic leukemia. Blood 121, 5068–5077. 10.1182/blood-2012-10-460170.

50. Wang, J., Liu, Y., Li, Z., Du, J., Ryu, M.J., Taylor, P.R., Fleming, M.D., Young, K.H., Pitot, H., and Zhang, J. (2010). Endogenous oncogenic Nras mutation promotes aberrant GM-CSF signaling in granulocytic/monocytic precursors in a murine model of chronic myelomonocytic leukemia. Blood 116, 5991–6002. 10.1182/blood-2010-04-281527.

51. Binder, M., Carr, R.M., Lasho, T.L., Finke, C.M., Mangaonkar, A.A., Pin, C.L., Berger, K.R., Mazzone, A., Potluri, S., Ordog, T., et al. (2022). Oncogenic gene expression and epigenetic remodeling of cis-regulatory elements in ASXL1-mutant chronic myelomonocytic leukemia. Nature Communications 13, 1434. 10.1038/s41467-022-29142-6.

52. Yoshimi, A., Balasis, M.E., Vedder, A., Feldman, K., Ma, Y., Zhang, H., Lee, S.C., Letson, C., Niyongere, S., Lu, S.X., et al. (2017). Robust patient-derived xenografts of MDS/MPN overlap syndromes capture the unique characteristics of CMML and JMML. Blood 130, 397–407. 10.1182/blood-2017-01-763219.

53. Gbyli, R., Song, Y., Liu, W., Gao, Y., Biancon, G., Chandhok, N.S., Wang, X., Fu, X., Patel, A., Sundaram, R., et al. (2022). In vivo anti-tumor effect of PARP inhibition in IDH1/2 mutant MDS/AML resistant to targeted inhibitors of mutant IDH1/2. Leukemia 36, 1313–1323. 10.1038/s41375-022-01536-x.

54. Krause, D.S., Fulzele, K., Catic, A., Sun, C.C., Dombkowski, D., Hurley, M.P., Lezeau, S., Attar, E., Wu, J.Y., Lin, H.Y., et al. (2013). Differential regulation of myeloid leukemias by the bone marrow microenvironment. Nature Medicine 19, 1513–1517. 10.1038/nm.3364.

55. Lewis, A.C., Pope, V.S., Tea, M.N., Li, M., Nwosu, G.O., Nguyen, T.M., Wallington-Beddoe, C.T., Moretti, P.A.B., Anderson, D., Creek, D.J., et al. (2022). Ceramide-induced integrated stress response overcomes Bcl-2 inhibitor resistance in acute myeloid leukemia. Blood 139, 3737–3751. 10.1182/blood.2021013277.

56. Patnaik, M.M., and Tefferi, A. (2022). Chronic myelomonocytic leukemia: 2022 update on diagnosis, risk stratification, and management. American Journal of Hematology 97, 352–372. https://doi.org/10.1002/ajh.26455.

57. Kim, K.-T., Baird, K., Ahn, J.-Y., Meltzer, P., Lilly, M., Levis, M., and Small, D. (2005). Pim-1 is up-regulated by constitutively activated FLT3 and plays a role in FLT3-mediated cell survival. Blood 105, 1759–1767. https://doi.org/10.1182/blood-2004-05-2006.

58. Mizuki, M., Schwäble, J., Steur, C., Choudhary, C., Agrawal, S., Sargin, B., Steffen, B., Matsumura, I., Kanakura, Y., Böhmer, F.D., et al. (2003). Suppression of myeloid transcription factors and induction of STAT response genes by AML-specific Flt3 mutations. Blood 101, 3164–3173. https://doi.org/10.1182/blood-2002-06-1677.

59. Chen, W., Kumar, A.R., Hudson, W.A., Li, Q., Wu, B., Staggs, R.A., Lund, E.A., Sam, T.N., and Kersey, J.H. (2008). Malignant Transformation Initiated by <em>Mll-AF9</em>: Gene Dosage and Critical Target Cells. Cancer Cell 13, 432–440. 10.1016/j.ccr.2008.03.005.

60. You, X., Wen, Z., Kong, G., Rajagopalan, A., Ranheim, E.A., Zhou, Y., Patnaik, M.M., and Zhang, J. (2019). Loss of Asxl1 Cooperates with Oncogenic Nras to Drive CMML Progression. Blood 134, 3790–3790. 10.1182/blood-2019-131347.

61. Fiskus, W., Sharma, S., Qi, J., Shah, B., Devaraj, S.G.T., Leveque, C., Portier, B.P., Iyer, S., Bradner, J.E., and Bhalla, K.N. (2014). BET Protein Antagonist JQ1 Is Synergistically Lethal with FLT3 Tyrosine Kinase Inhibitor (TKI) and Overcomes Resistance to FLT3-TKI in AML Cells Expressing FLT-ITD. Molecular Cancer Therapeutics 13, 2315–2327. 10.1158/1535-7163.Mct-14-0258.

62. Buonamici, S., Yoshimi, A., Thomas, M., Seiler, M., Chan, B., Caleb, B., Darman, R., Fekkes, P., Karr, C., Keaney, G.F., et al. (2016). H3B-8800, an Orally Bioavailable Modulator of the SF3b Complex, Shows Efficacy in Spliceosome-Mutant Myeloid Malignancies. Blood 128, 966–966. 10.1182/blood.V128.22.966.966.

63. Jang, J.E., Eom, J.-I., Jeung, H.-K., Cheong, J.-W., Lee, J.Y., Kim, J.S., and Min, Y.H. (2017). Targeting AMPK-ULK1-mediated autophagy for combating BET inhibitor resistance in acute myeloid leukemia stem cells. Autophagy 13, 761–762. 10.1080/15548627.2016.1278328.

64. Saenz, D.T., Fiskus, W., Manshouri, T., Rajapakshe, K., Krieger, S., Sun, B., Mill, C.P., DiNardo, C., Pemmaraju, N., Kadia, T., et al. (2017). BET protein bromodomain inhibitor-based combinations are highly active against post-myeloproliferative neoplasm secondary AML cells. Leukemia 31, 678–687. 10.1038/leu.2016.260.

65. Qian, K.C., Studts, J., Wang, L., Barringer, K., Kronkaitis, A., Peng, C., Baptiste, A., LaFrance, R., Mische, S., and Farmer, B. (2005). Expression, purification, crystallization and preliminary crystallographic analysis of human Pim-1 kinase. Acta Crystallographica Section F 61, 96–99. doi:10.1107/S1744309104029963.

66. Karatas, O.F. (2018). Antiproliferative potential of miR-33a in laryngeal cancer Hep-2 cells via targeting PIM1. Head Neck 40, 2455–2461. 10.1002/hed.25361.

67. Wang, Y., Zhou, X., Shan, B., Han, J., Wang, F., Fan, X., Lv, Y., Chang, L., and Liu, W. (2015). Downregulation of microRNAIZl33a promotes cyclinIZldependent kinase 6, cyclin D1 and PIM1 expression and gastric cancer cell proliferation. Mol Med Rep 12, 6491–6500. 10.3892/mmr.2015.4296.

68. Ianevski, A., Giri, A.K., and Aittokallio, T. (2020). SynergyFinder 2.0: visual analytics of multi-drug combination synergies. Nucleic Acids Research 48, W488–W493. 10.1093/nar/gkaa216.

69. Dobin, A., Davis, C.A., Schlesinger, F., Drenkow, J., Zaleski, C., Jha, S., Batut, P., Chaisson, M., and Gingeras, T.R. (2013). STAR: ultrafast universal RNA-seq aligner. Bioinformatics 29, 15–21. 10.1093/bioinformatics/bts635.

70. Love, M.I., Huber, W., and Anders, S. (2014). Moderated estimation of fold change and dispersion for RNA-seq data with DESeq2. Genome Biology 15, 550. 10.1186/s13059-014-0550-8.

71. Subramanian, A., Tamayo, P., Mootha, V.K., Mukherjee, S., Ebert, B.L., Gillette, M.A., Paulovich, A., Pomeroy, S.L., Golub, T.R., Lander, E.S., and Mesirov, J.P. (2005). Gene set enrichment analysis: A knowledge-based approach for interpreting genome-wide expression profiles. Proceedings of the National Academy of Sciences 102, 15545–15550. doi:10.1073/pnas.0506580102.

