## Supplemental Figures for "Targeting BET Proteins downregulates miR-33a to promote synergy with PIM inhibitors in CMML"

**Supplemental Figure S1.**

A.

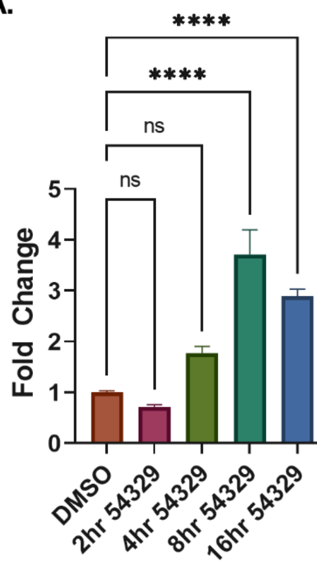


B.

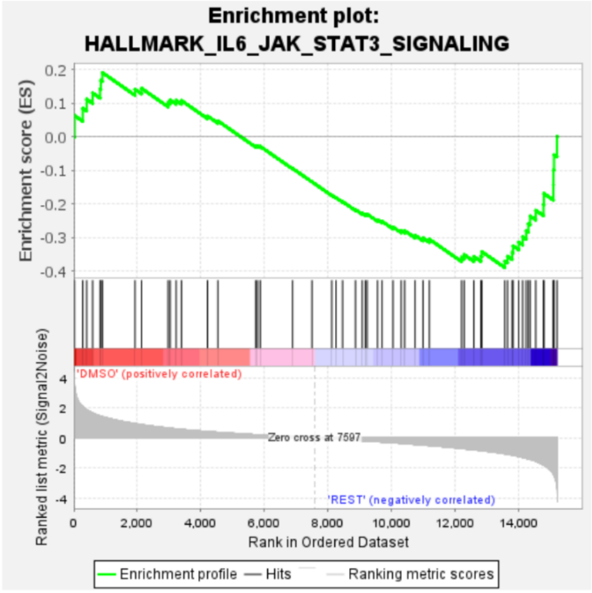


C.

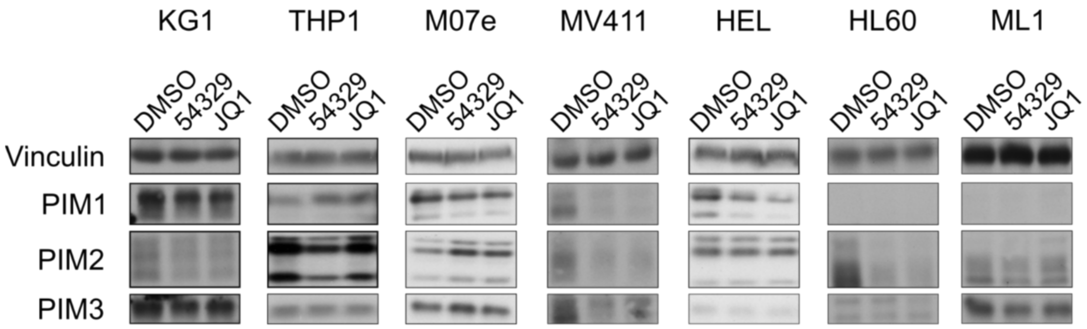

**Supplemental Figure S2.**

A.

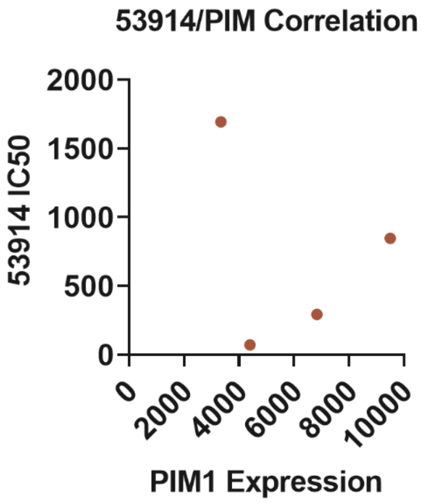

**Supplemental Figure S3.**

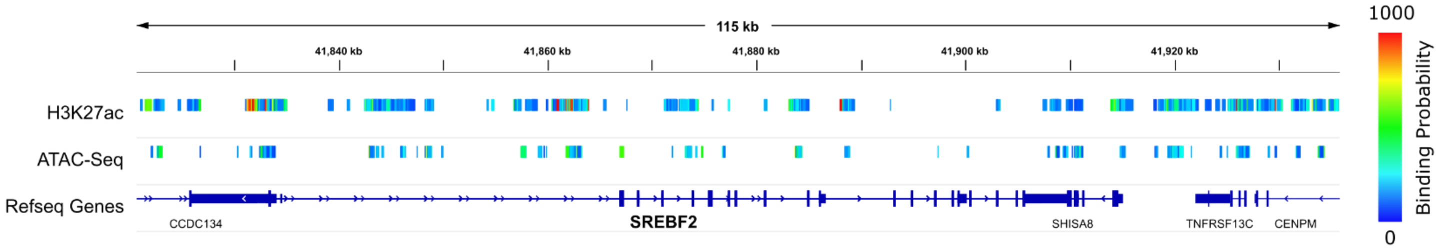

**Supplemental Figure S4.**

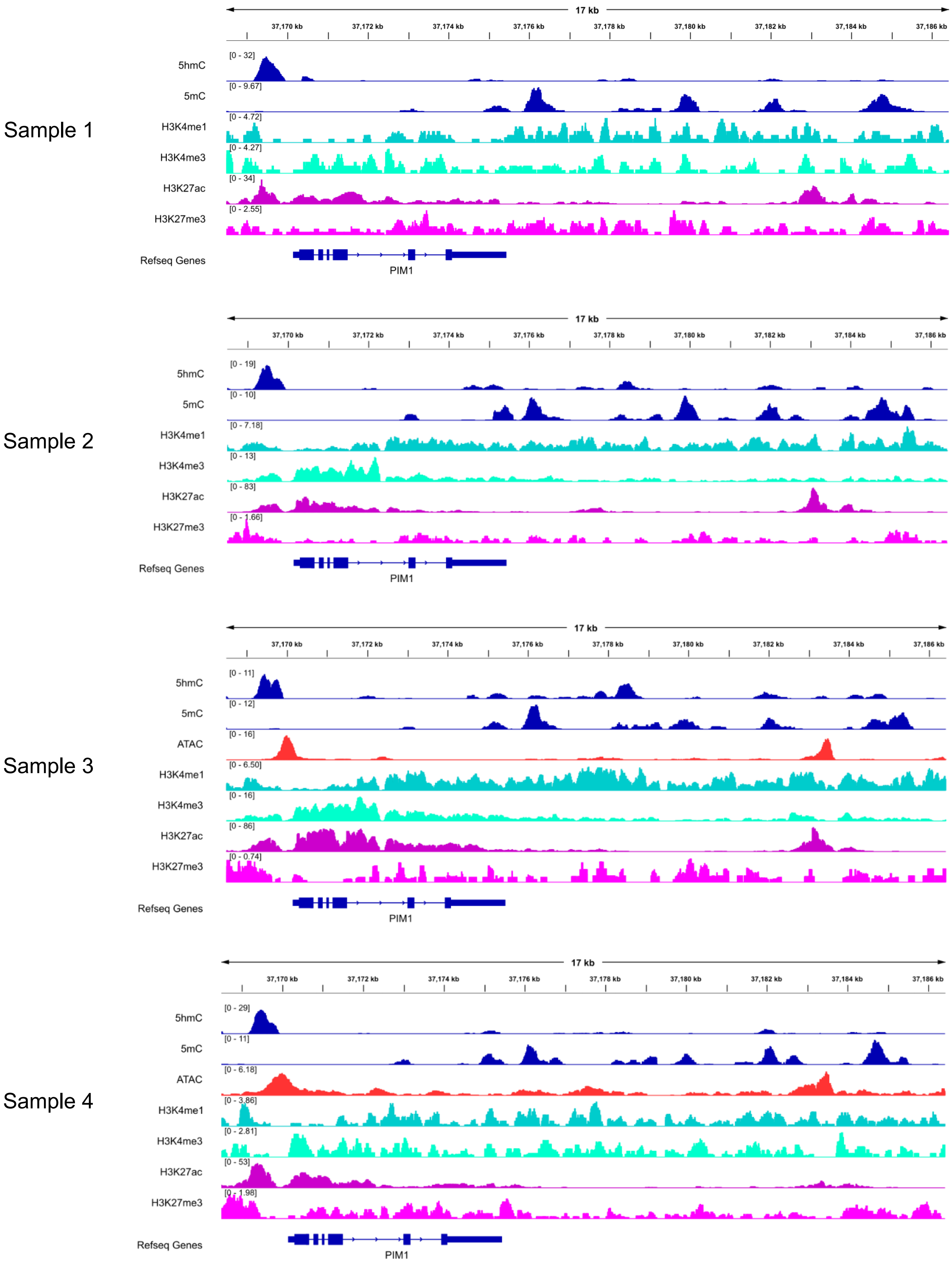

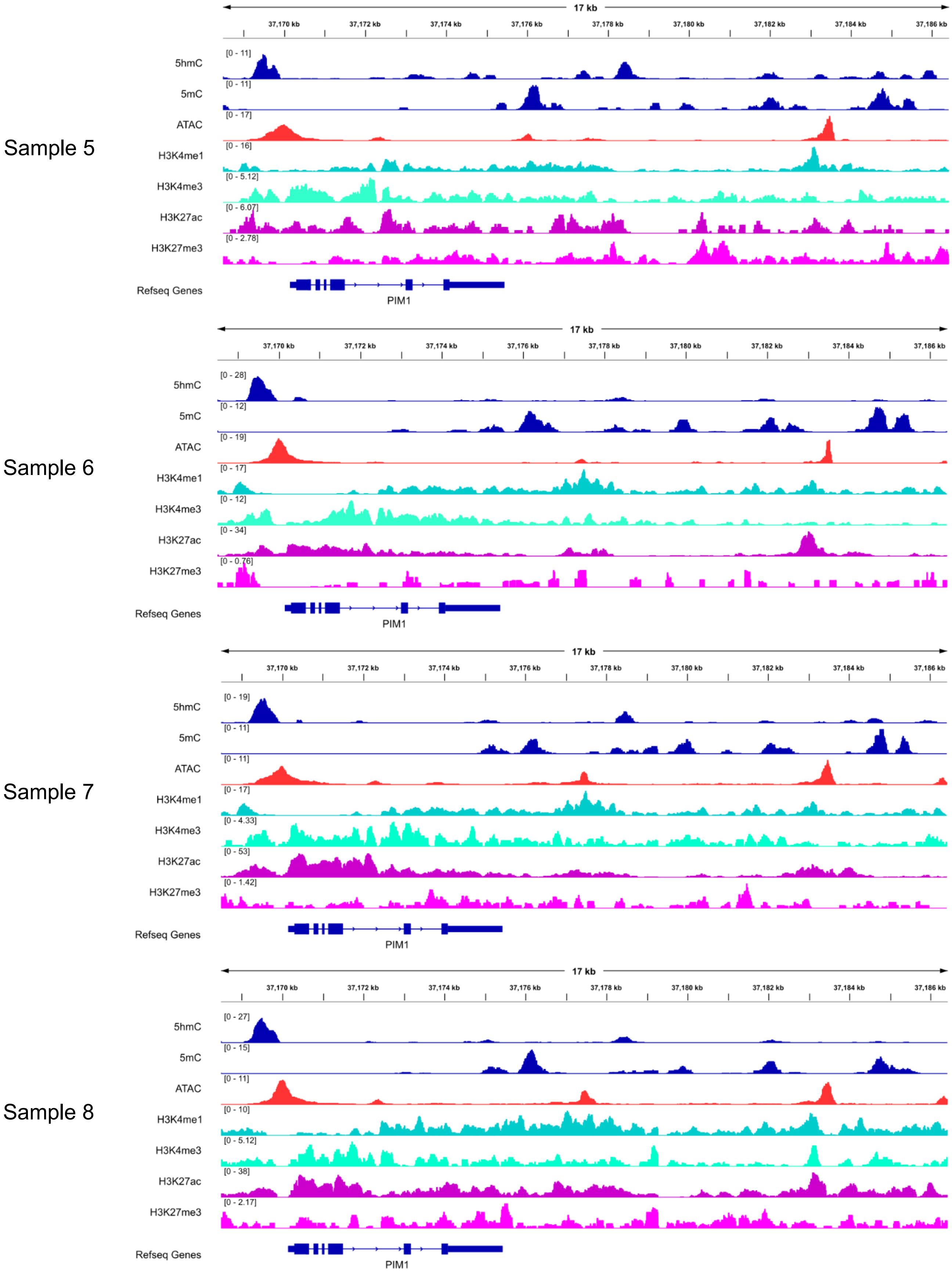

Sample 5
17 kb
37,170 kb
37,172 kb
37,174 kb
37,176 kb
37,178 kb
37,180 kb
37,182 kb
37,184 kb
37,186 kb
5hmC [0 - 11]
5mC [0 - 11]
ATAC [0 - 17]
H3K4me1 [0 - 16]
H3K4me3 [0 - 5.12]
H3K27ac [0 - 6.07]
H3K27me3 [0 - 2.78]
Refseq Genes
PIM1
Sample 6
17 kb
37,170 kb
37,172 kb
37,174 kb
37,176 kb
37,178 kb
37,180 kb
37,182 kb
37,184 kb
37,186 kb
5hmC [0 - 28]
5mC [0 - 12]
ATAC [0 - 19]
H3K4me1 [0 - 17]
H3K4me3 [0 - 12]
H3K27ac [0 - 34]
H3K27me3 [0 - 0.76]
Refseq Genes
PIM1
Sample 7
17 kb
37,170 kb
37,172 kb
37,174 kb
37,176 kb
37,178 kb
37,180 kb
37,182 kb
37,184 kb
37,186 kb
5hmC [0 - 19]
5mC [0 - 11]
ATAC [0 - 11]
H3K4me1 [0 - 17]
H3K4me3 [0 - 4.33]
H3K27ac [0 - 53]
H3K27me3 [0 - 1.42]
Refseq Genes
PIM1
Sample 8
17 kb
37,170 kb
37,172 kb
37,174 kb
37,176 kb
37,178 kb
37,180 kb
37,182 kb
37,184 kb
37,186 kb
5hmC [0 - 27]
5mC [0 - 15]
ATAC [0 - 11]
H3K4me1 [0 - 10]
H3K4me3 [0 - 5.12]
H3K27ac [0 - 38]
H3K27me3 [0 - 2.17]
Refseq Genes
PIM1

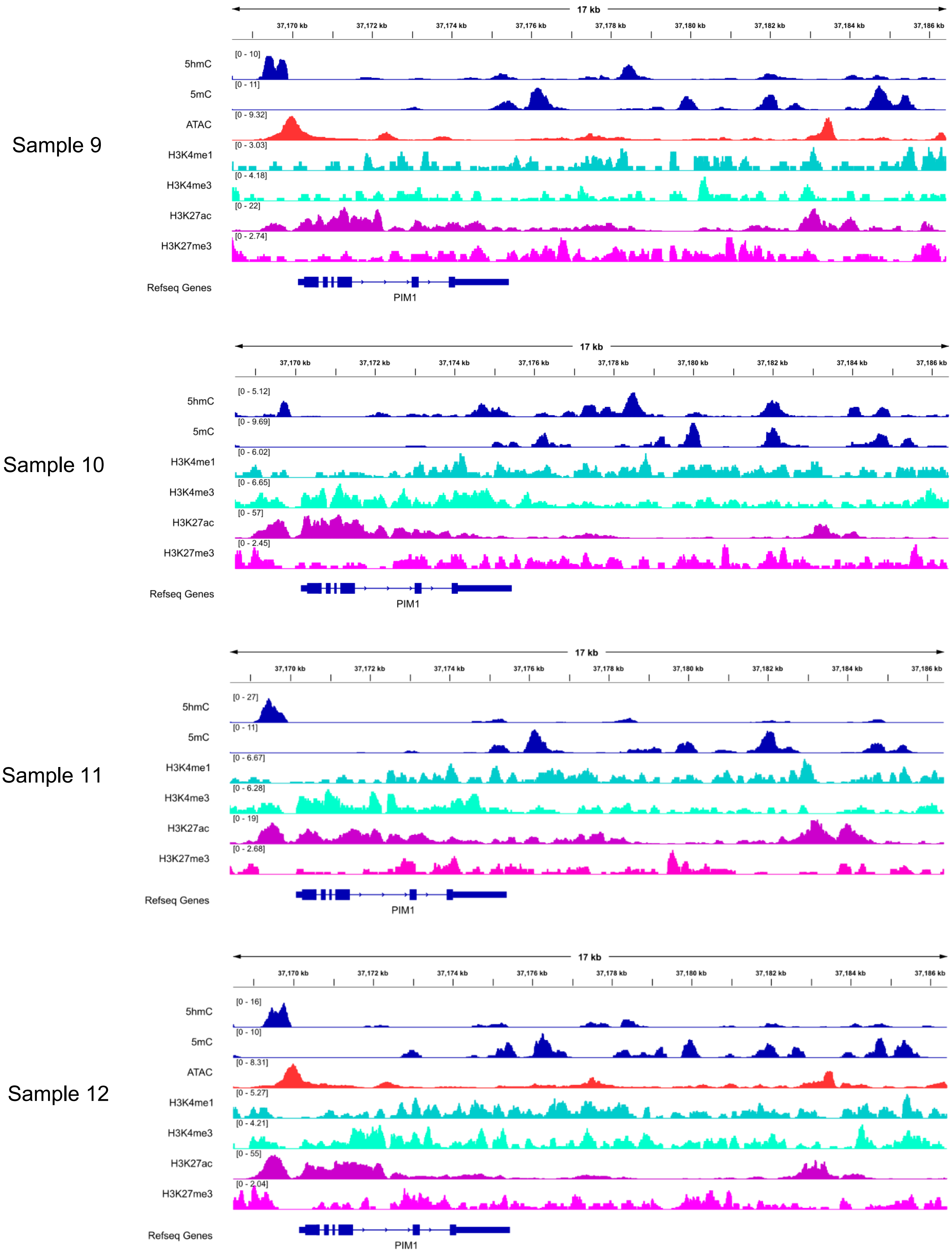

Sample 9
17 kb
37,170 kb
37,172 kb
37,174 kb
37,176 kb
37,178 kb
37,180 kb
37,182 kb
37,184 kb
37,186 kb
5hmC [0 - 10]
5mC [0 - 11]
ATAC [0 - 9.32]
H3K4me1 [0 - 3.03]
H3K4me3 [0 - 4.18]
H3K27ac [0 - 22]
H3K27me3 [0 - 2.74]
Refseq Genes
PIM1
Sample 10
17 kb
5hmC [0 - 5.12]
5mC [0 - 9.69]
H3K4me1 [0 - 6.02]
H3K4me3 [0 - 6.65]
H3K27ac [0 - 57]
H3K27me3 [0 - 2.45]
Refseq Genes
PIM1
Sample 11
17 kb
5hmC [0 - 27]
5mC [0 - 11]
H3K4me1 [0 - 6.67]
H3K4me3 [0 - 6.28]
H3K27ac [0 - 19]
H3K27me3 [0 - 2.68]
Refseq Genes
PIM1
Sample 12
17 kb
5hmC [0 - 16]
5mC [0 - 10]
ATAC [0 - 8.31]
H3K4me1 [0 - 5.27]
H3K4me3 [0 - 4.21]
H3K27ac [0 - 55]
H3K27me3 [0 - 2.04]
Refseq Genes
PIM1

Sample 13

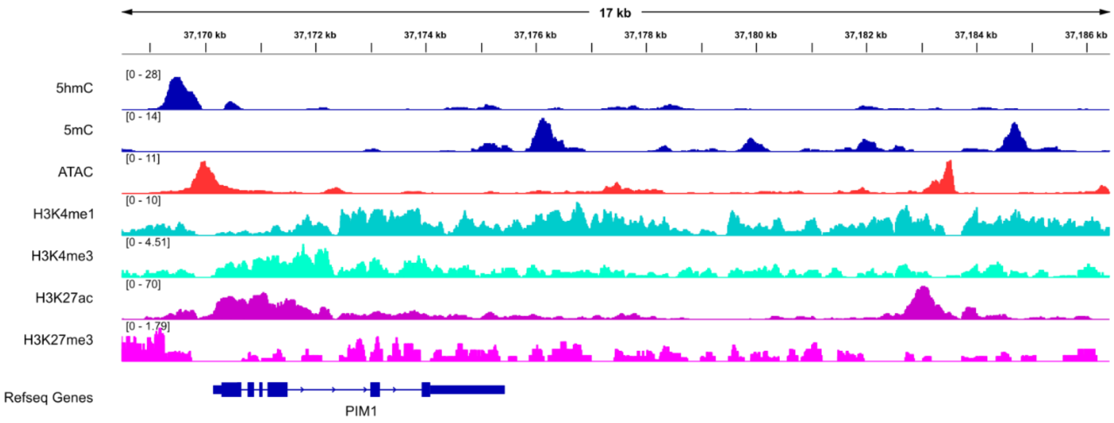


Sample 14

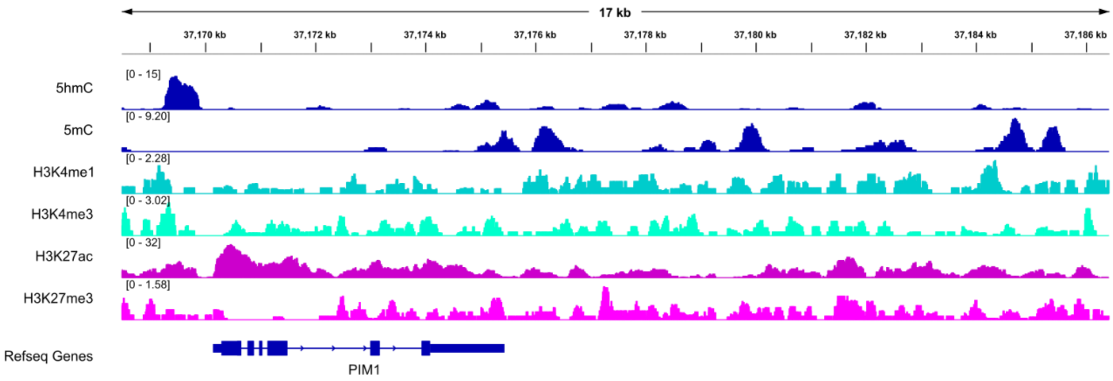


Sample 15

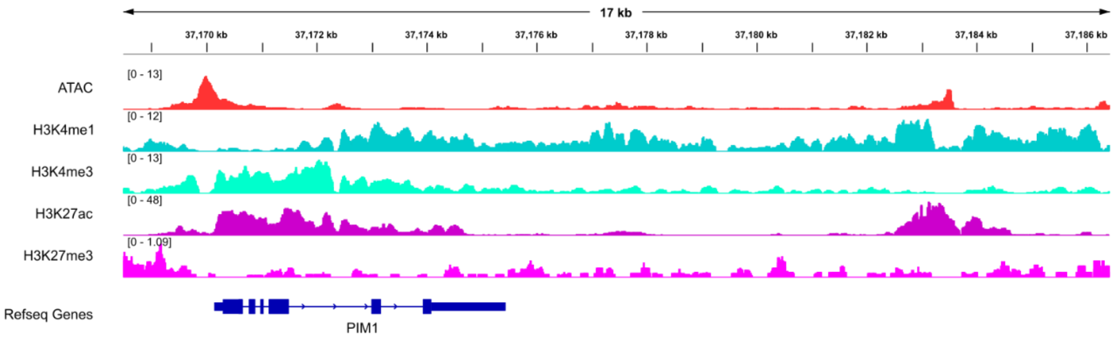


Sample 16

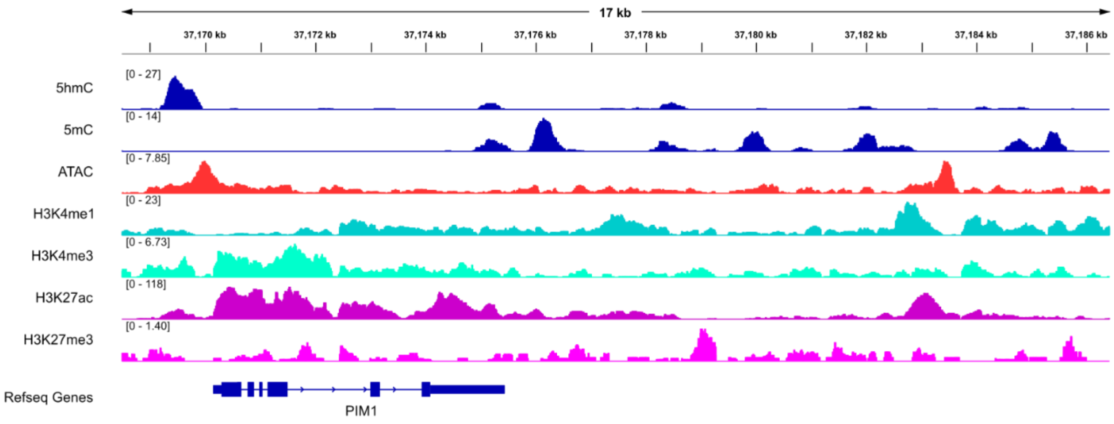

**Supplemental Figure S5.**

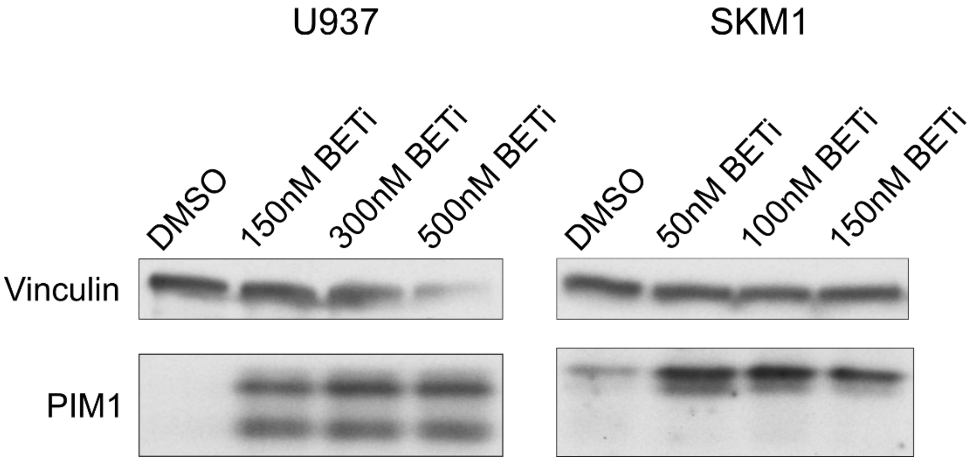
