## Supplemental Table 3 for "Targeting BET Proteins downregulates miR-33a to promote synergy with PIM inhibitors in CMML"

| Patient samples used in PDX | | | | | | | |
| --- | --- | --- | --- | --- | --- | --- | --- |
| ID | AGE | SEX | WHO | FAB | MAYO | Blast% | Mutations |
| RX-1-001 | unknown | unknown | CMML-0 | MPN-CMML | 2 | 1 | SRSFF2 (c.43T>G, p.S15A; 50.8%), NRAS (c.35G>A, p.G12D; 32.7%), TET2 (c.5037C>A, p.Y1679*; 48.8%), TET2 (c.2305delC, p. Q769Sfs*44, 39%) |
| 3-F-004 | 67 | Male | CMML-2 | MD-CMML | 2 | 16 | NRAS (p.G12S), SF3B1 (p.K666N), TET2 (p.R123fs and p.H839fs) |
| 3-G-004 | 60 | Female | CMML-0 | MP-CMML | 2 | 1 | TET2, CBL, SRSF2, SETBP1 |
| KB-10-005 | 75 | Female | unknown | unknown | unknown | unknown | NRAS, ASXL1, TET2, ZRSR2 |
| Patient samples used in CFA | | | | | | | |
| 2-C-001 | 71 | Male | CMML-1 | MPN-CMML | unknown | 5 | TET2, SRSF2, NRAS |
| 2-J-001 | 57 | female | unknown | unknown | unknown | unknown | KRAS, NOTCH1 |
| 2-V-001 | 76 | Male | CMML-0 | MP-CMML | 3 | 5.5 | TET2(p.G1703* C.5107G>T, p.N1823Kfs*23), CBL(p.P417R), SRSF2(p.P95L) |
| 3-D-001 | 71 | Male | CMML-0 | MP-CMML | unknown | unknown | CEBPA, EZH2, NRAS, STAG2, TET2 |
| 3-E-002 | 54 | Male | CMML-1 | MP-CMML | 3 | 5 | SRSF2(p.P95H), TET2(p.Q734Rfs*17, c.2198del) |
| 3-G-001 | 60 | Female | CMML-0 | MPN-CMML | 2 | 1 | TET2, CBL, SRSF2, SETBP1 |
| 2-D-001 | 82 | Male | CMML-2 | MP-CMML | 2 | 5 | NRAS, SRSF2, SETBP1, ASXL1 |
| 3-F-001 | 67 | Male | CMML-2 | MD-CMML | 2 | 15 | TET2, NRAS, SF3B1 |
| 1-A-001 | 49 | female | CMML-0 | MD-CMML | 3 | 1.2 | CEBPA, KRAS |
| 2-U-001 | 56 | Female | CMML-0 | MPN-CMML | 3 | 4 | from 8 months prior: TET2 (c.5735A>G, p.H1912R), TET2 (c.2491C>T, p. Q831*), SETBP1 (c. 1879C>T, p. R627C), NRAS (c. 35G>T, p. G12V) |

**Supplemental Table 3.** Characteristics of Patients used in PDX and CFA Experiments.
